# Skin capillary endothelial cells form a network of spatiotemporally conserved Ca^2+^ activity

**DOI:** 10.1101/2025.08.15.669933

**Authors:** Anush Swaminathan, David G. Gonzalez, Catherine Matte-Martone, Fei Xu, Deandra Simpson, Jessica L. Moore, Zhongqi Lin, Ushnish Rana, David Monedero-Alonso, Julia J. Mack, Chen Yuan Kam, Valentina Greco

**Author notes:** Co-correspondence to: Valentina Greco, Chen Yuan Kam, Julia J. Mack.

## Abstract

Ca^2+^ signaling and its regulation are important for endothelial cell (EC) function and signaling. Yet, the spatiotemporal organization of Ca^2+^ activity and its regulation across a vascular plexus is poorly understood in an *in vivo* mammalian context. To overcome this gap in knowledge, we developed an intravital imaging approach to resolve Ca^2+^ activity with single cell resolution in skin vasculature of adult mice via multiphoton microscopy. Here, we tracked thousands of Ca^2+^ events in the skin capillary plexus during homeostasis and observed signaling heterogeneity between ECs, with just over half displaying Ca^2+^ activity at any given time. Longitudinal tracking of the same mice revealed that the same capillary ECs maintain Ca^2+^ activity over days to weeks. Interestingly, activity dynamics, such as frequency and event duration, are not conserved at a single cell level but are maintained at an EC population level. Molecularly, conditional deletion of the gap junction protein Connexin 43 (Cx43cKO) in ECs lead to a subset of ECs displaying sustained Ca^2+^ activity, biasing signaling dynamics of the whole network towards chronically persistent activity over time. Sustained capillary Ca^2+^ activity resulted in vascular permeability and flow dysregulation. Lastly, through pharmacological targeting of known agonists/antagonists, we showed that inhibition of L-type Voltage Gated Ca^2+^ channels (VGCCs) non cell-autonomously restores Ca^2+^ activity, blood flow, and barrier function in Cx43cKO mice. Collectively, our work provides insight into the characteristics, extent, and regulation of Ca^2+^ activity in skin capillaries of live mice with unprecedented spatial and temporal resolution.

**Significance Statement:** Ca^2+^ signaling in mammalian endothelial cells (ECs) locally regulates blood flow, force sensing, and vessel permeability. Past studies have investigated Ca^2+^ signaling during vascular remodeling and repair. However, there is a gap in our understanding of how tissue-level Ca^2+^ is spatiotemporally organized and regulated during homeostasis. Intravital imaging in skin vasculature of live mice reveals that a conserved network of ECs participates in tissue-wide Ca^2+^ signaling over weeks. How this network maintains itself over time requires cellular communication through gap junction protein Connexin 43 (Cx43). Loss of EC Cx43 leads to heightened plexus-wide Ca^2+^ activity, and vessel barrier and flow dysregulation. Inhibition of L-type Ca^2+^ channels non-cell autonomously restores the capillary Ca^2+^ landscape, and rescues both barrier and flow dysfunction.

## Introduction

Ca^2+^ is an essential second messenger, mediating tightly regulated signaling pathways to integrate spatial and temporal information across multiple scales, from individual cell transients to multi-cell patterns (1–5). *In vitro*, the temporal dynamics of Ca^2+^ activity within cells differentially encodes downstream functions, enabling its versatility as a signaling pathway (6–11). In excitable tissues *in vivo*, such as basal and lateral amygdala (BLA) after application of a conditioning stimuli, there is considerable overlap in which specific neurons are active over days (12). Moreover, the BLA network displays tightly conserved population-level properties, such as maintenance of total proportion of Ca^2+^ active neurons over time (12). In non-excitable tissues *in vivo*, we previously demonstrated that cells in the epidermal stem cell compartment engage in both local and large-scale Ca^2+^ activity without any stimuli (4). These tissue-wide analyses revealed that all epidermal stem cells display Ca^2+^ activity within one day during homeostasis. However, *in vivo* Ca^2+^ activity, characteristics, and regulation, across other tissues, especially in mammalian systems, remains relatively unexplored.

Endothelial cells (ECs) are specialized squamous cells that line the lumen of all blood vessels (13). The endothelium regulates key functions in the vasculature through Ca^2+^ activity, such as angiogenesis, vessel tone, local blood flow control, mechanotransduction, and barrier integrity (14–19). *In vivo* work in zebrafish vasculature during development has tracked Ca^2+^ across EC populations and identified spatiotemporally organized Ca^2+^ activity specifically in cells driving angiogenesis (tip cells) (20). These cells also display variable Ca^2+^ temporal dynamics which shape downstream vessel fates (20). Recently, more studies have focused on mouse brain capillaries given their extensive neurovascular coupling and outsized energy and metabolic demands, to identify molecular mechanisms driving Ca^2+^ signaling within individual ECs. These include IP_3_R-mediated Ca^2+^ release, and how channels including Kir2.1, TRPV4, TRPA1, and Piezo1 shape Ca^2+^ influx (21–25). Yet we lack a spatiotemporal understanding of Ca^2+^ signaling dynamics during homeostasis across more general, peripheral vascular plexi in their native, unperturbed environment.

Organization of Ca^2+^ activity also involves information exchange between cells, as ECs can coordinate Ca^2+^ activity in response to different homeostatic cues (26–30). While individual ECs can regulate cytoplasmic Ca^2+^ concentrations through a variety of mechanisms, multicellular coordination of Ca^2+^ activity between ECs has largely been studied in the context of gap junction-mediated intercellular communication (31). Gap junctions are intercellular channels that allow for cell-cell communication in all tissues, including the vasculature, through electrical coupling and transfer of second messengers such as IP_3_ (32, 33). Gap junctions in the vascular endothelium can facilitate fast, local propagation of vasodilatory signals and enable efficient neurovascular coupling (26, 34–36). Additionally, gap junction activity can modulate EC function during homeostasis, inflammation, and injury, through ATP release and purinergic signaling pathways (37–39). Evidence for EC coordination during homeostasis, development, and injury indicates that molecular signaling may not only be influenced by direct neighbors but also regulated on a tissue-wide scale (40–44). Gap junctions have been implicated in coordination of Ca^2+^ signaling networks in other tissues, such as the loss of Connexin 43 (Cx43) gap junctions leading to tissue-wide uncoupling of coordination across the epidermal stem cell compartment and repeated, more frequent, and longer signaling within restricted neighborhoods (4). In *Drosophila* lymph glands, inhibiting gap junctions alters the Ca^2+^ signaling frequency and promotes precocious differentiation of blood progenitors (45).

Despite these important insights, it remains difficult to study Ca^2+^ dynamics in an *in vivo* mammalian system given the technical challenges of tracking ECs and their signaling across space and time. Here, we overcome this challenge by focusing on the skin capillary plexus, due to its critical roles in nutrient and oxygen delivery, and tracking Ca^2+^ dynamics in hundreds of ECs over time through our non-invasive, intravital imaging approach (Figure S1) (4, 42, 46). We observed extensive Ca^2+^ activity across capillary plexus ECs during homeostasis, with patterns of Ca^2+^ activity conserved spatially within the same cells and temporally at the network level. To probe how the spatiotemporal organization of EC Ca^2+^ activity is coordinated, we conditionally deleted Connexin 43 (Cx43cKO) in ECs, and discovered chronically sustained Ca^2+^ activity, altered blood flow dynamics, barrier dysfunction, and dysregulation of EC temporal coordination across the capillary plexus. Finally, we show that inhibition of L-type Voltage Gated Ca^2+^ channels (VGCCs) can restore Ca^2+^ activity in Cx43cKO ECs, along with blood flow and barrier function. Altogether, this study defines the spatiotemporal organization and regulation of Ca^2+^ activity by a conserved EC network in the skin capillary plexus.

## Results

### Spatiotemporal analyses of the skin capillary plexus reveal a network of endothelial cells with conserved Ca^2+^ activity over time

Understanding plexus-wide Ca^2+^ activity is fundamental to reveal how ECs coordinate their signaling to achieve proper vascular function. The superficial nature of the skin capillary plexus makes it especially tractable to longitudinal tracking of an *in vivo* EC population through non-invasive multi-photon microscopy. Here, we aimed to gain a plexus-wide view of Ca^2+^ activity in skin capillaries with single-cell resolution. Towards this goal, we recorded Ca^2+^ activity with simultaneous imaging of Ca^2+^ dynamics and EC nuclei via genetically encoded fluorescent reporters. We used an EC-specific inducible Cre driver under the control of the vascular endothelial cadherin (VECad) promoter to simultaneously recombine a nuclear H2B-mCherry reporter and GCaMP6s, a Ca^2+^ reporter encoding a calmodulin-GFP fusion protein (VECadCreER; Rosa26-CAG-LSL-GCaMP6s; LSL-H2B-mCherry mice induced with 2 mg Tamoxifen daily for 4 consecutive days; Figure S1 A). We focused on a simplified skin model devoid of hair follicles, palmoplantar skin, and recorded Ca^2+^ activity in the entire capillary plexus (4 to 5 regions per mouse, with 100-150 cells per region across 12 microns in depth and 5 optical sections) over a period of 17 minutes and 12 seconds (300 frames at 3.44 seconds/frame) in adult (2-4 months old) anesthetized mice via two-photon microscopy (Figure S1). During these timelapse recordings, we observed dynamic and heterogeneous Ca^2+^ activity across the plexus with hundreds of events per region and different levels of Ca^2+^ activity and dynamics between cells (Figure 1 A; Movie 1). To quantitatively analyze the dynamics of the observed Ca^2+^ activity, we devised a computational analysis pipeline that segments individual ECs by using the position of the nuclear H2B-mCherry signal as a proxy for the cell body (Figure S1 A-C). This enabled us to isolate and analyze changes in GCaMP intensity of each individual cell within the imaging region for the entire duration of the timelapse recording (Figure S1 C). The pipeline determines the number and duration of Ca^2+^ events per cell without being confounded by variable baseline Ca^2+^ levels, through counting any activity above a set threshold as an event (>50% change in average MFI above the minimum MFI) (Figure 1 A; Figure S1 C). We analyzed 1,765 cells across 4 mice and discovered that just over half of ECs (52.4 ± 7.34%) exhibited Ca^2+^ activity during the timeframe of recording (defined as having an ‘active’ cell status, with the others assigned a status of ‘inactive’; Figure 1 B).

**Figure 1.**
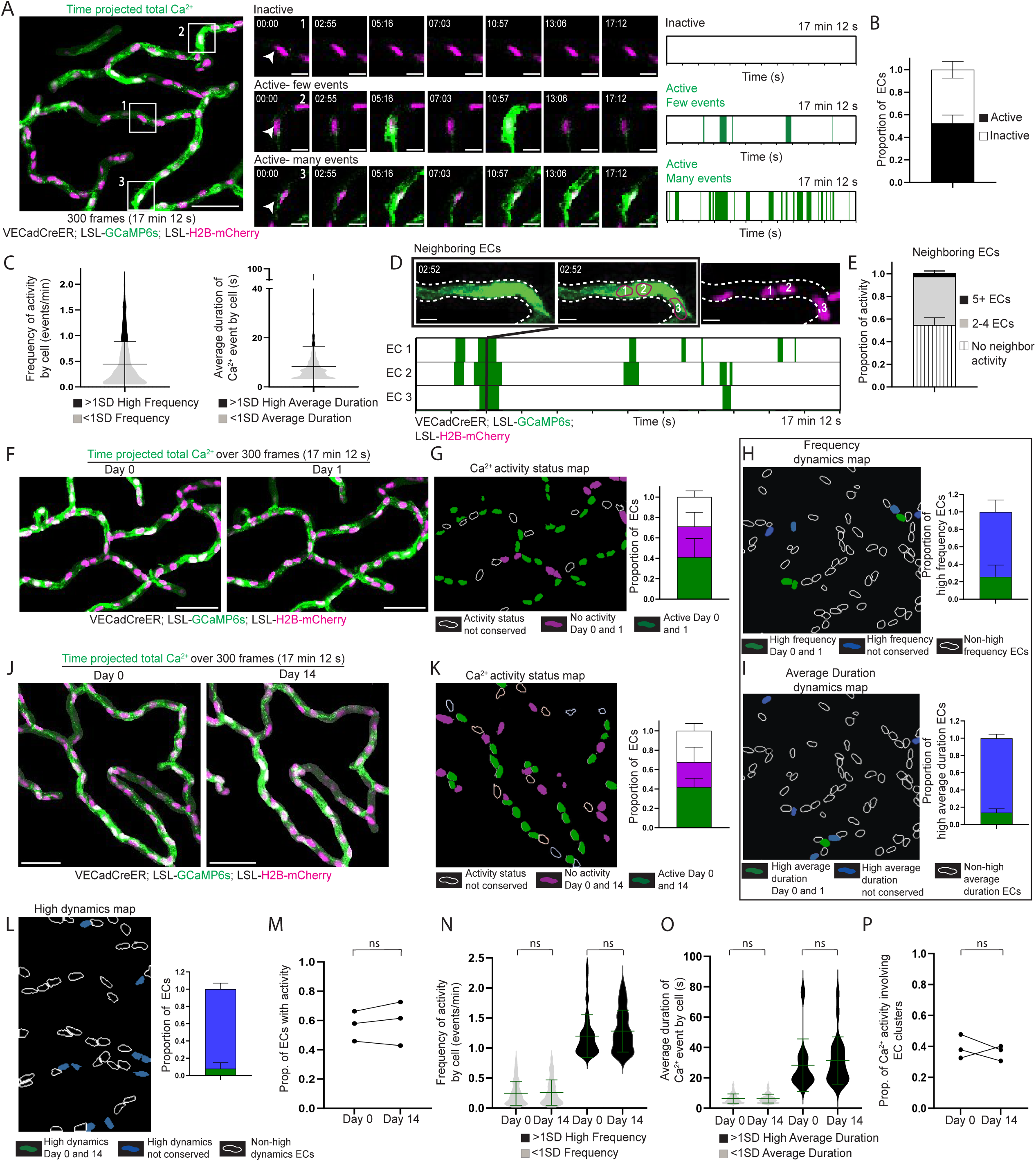
The skin capillary plexus displays Ca^2+^ activity orchestrated by a spatiotemporally conserved network of endothelial cells. **(A)** *Left*: Max intensity projection of GCaMP6s signal (green) with H2B-mCherry signal (magenta) from 300 frame (17 minutes 12 seconds) recording of skin capillary ECs showing heterogenous Ca^2+^ activity between cells (scale bar: 50 µm). *Mid*: Insets of 3 different regions represent differences in Ca^2+^ activity between cells (scale bar: 10 µm). *Right*: Ca^2+^ events and their durations over the recording time for each of the ECs displayed in the insets (green represents Ca^2+^ activity) **(B)** Proportion of ECs displaying Ca^2+^ activity (active ECs in black and inactive ECs in white). *n=*14 regions from 4 mice. **(C)** Frequency of Ca^2+^ activity per cell (events/min) and average duration (s) of a Ca^2+^ event per cell across 1,765 ECs from 4 mice. EC groups represented by black (>1SD high activity) and grey (<1SD activity) **(D)** *Top*: Ca^2+^ activity occurring simultaneously across 3 ECs (numbered and drawn in magenta outline matching H2B-mCherry) (scale bar: 10 µm). *Bottom*: Ca^2+^ events and their durations over recording time for each EC. Black line across plots indicates the timepoint when all 3 ECs simultaneously display Ca^2+^ activity. **(E)** Proportion of Ca^2+^ activity involving various EC cluster sizes. *n* = 14 regions from 4 mice. **(F)** Max intensity projection of GCaMP6s signal with H2B-mCherry signal, with the same regions revisited and recorded 24 hours later (scale bar: 50 µm). **(G)** *Left*: Ca^2+^ activity status map for ECs revisited after 24 hours in (F). Non-conserved activity status (white outline), inactive both days (magenta) and active both days (green). *Right*: Proportion of ECs by conservation of their activity status. Chi-square analysis (observed activity 24 hours after versus random activity as null hypothesis) P<0.0001. *n=*7 regions from 3 mice. **(H)** and **(I)** *Left*: Dynamics map for ECs revisited after 24 hours for conserved high frequency or high average duration behavior on Day 0 and Day 1 (green), not conserved (blue) or not observed (white outline) groups. *Right*: Conservation of >1SD dynamics behavior (green) as a proportion of total high frequency or high average duration ECs. *n=*7 regions from 3 mice. **(J)** Max intensity projection with the same region revisited and recorded 14 days later (scale bar: 50 µm). **(K)** *Left*: Ca^2+^ activity status map for ECs revisited after 14 days in (J). *Right*: Conservation of activity status as a proportion of total ECs. Chi-square (observed activity 14 days after versus random activity as null hypothesis) P<0.0001. *n=*8 regions from 3 mice. **(L)** *Left*: Dynamics map for ECs revisited after 14 days for conserved >1SD high dynamics (including both high frequency and high average duration). *Right*: Conservation of >1SD high dynamics behavior (green) as a proportion of total high frequency and high average duration ECs. *n=*8 regions from 3 mice. **(M)** Proportion of active ECs from 3 mice on Day 0 and Day 14; ns: P>0.05, paired t-test, *n=*457 total cells from 3 mice. **(N)** and **(O)** Frequency of Ca^2+^ activity (events/min) and average duration (s) of Ca^2+^ events per cell on Day 0 and Day 14. Black (>1SD high activity) and grey (<1SD activity); ns: P>0.05, unpaired t-test, *n=*265 active cells from 3 mice. **(P)** Proportion of Ca^2+^ activity involving EC clusters from 3 mice on Day 0 and Day 14; ns: P>0.05, paired t-test.

Considering the GCaMP6s sensor decays with a half-time of approximately 0.6-0.9 seconds (47, 48), we wanted to test to what extent the chosen 3.44 seconds/frame imaging parameters may affect our ability to accurately capture EC Ca^2+^ dynamics. Comparing 3.44 seconds/frame imaging to 0.62 seconds/frame imaging (using reduced optical sections and at the expense of visualizing network connectivity) showed no significant differences in the percentage of signaling ECs captured (52.4 ± 7.34% at 3.44 seconds/frame vs. 62.8 ± 13.8% at 0.62 seconds/frame) and demonstrated similar spread in Ca^2+^ event durations (134.2 seconds maximum duration at 3.44 seconds/frame imaging vs. 132.7 seconds maximum duration at 0.62 seconds/frame imaging) (Figure S1 D-E). Given our goal was to interrogate plexus wide Ca^2+^ dynamics, we applied the 3.44 seconds/frame imaging parameter to capture the entire 3D capillary plexus (Figure S1 D) and quantify signaling activity. Using our pipeline, we found that the majority of active ECs display dynamics within a defined range in both frequency (0.445 ± 0.440 events per minute per cell, encompassing 83.9 ± 7.50% of cells) and average duration (8.36 ± 8.15 seconds per event per cell, encompassing 91.3 ± 3.84% of ECs) of Ca^2+^ events (Figure 1 C). A small population of ECs displayed activity above that range, indicating either high frequency (1.31 ± .351 events/minute) or high average duration (29.8 ± 16.4 seconds per event) dynamics (Figure 1 C; Figure S2 A). Lastly, to understand the prevalence of multi-cellular activity, we adapted our pipeline to identify neighboring cells displaying Ca^2+^ activity either within the same time frame, or one frame apart, and calculated the number of cells involved (Figure 1 D; Figure S1 A-B). We found that the most represented cluster size for EC Ca^2+^ activity occurred across 2-4 cells (42.8 ± 5.64% of total activity) with a small percentage involving 5 or more cells (2.97 ± 1.32% of total activity) (Figure 1 E). Comparing imaging frequency also showed a similar spread in EC cluster sizes (up to 12 cells at 3.44 seconds/frame imaging vs. 11 cells at 0.62 seconds/frame imaging) (Figure S1 E).

After we established a quantitative understanding of EC Ca^2+^ dynamics, we next asked how activity status is regulated over longer periods of time. Towards this goal, we leveraged our longitudinal imaging approach that allows us to revisit the same cells over days to weeks. We began with short-term revisits and intriguingly, discovered that not only was the proportion of active and inactive cells largely conserved after 24 hours (58.7 ±14.4% active on Day 0 vs. 52.0 ± 18.8% on Day 1), but activity status was also conserved at the cellular level (71.1 ± 6.03% of cells maintained their activity status after 24 hours) (Figures 1 F and G; Figure S2 B; Movie 2). We also observed that only 28.9 ± 7.89% of high frequency and 17.0 ± 6.30% of high average duration ECs retained their dynamics after 24 hours, indicating that despite maintaining activity status (active vs. inactive), individual ECs largely exhibit different Ca^2+^ dynamics (frequency and average duration) over days (Figure 1 H and I; Figure S2 C).

Next, we asked whether Ca^2+^ activity status is also conserved over a longer time frame. As our past work has demonstrated that ECs remain positionally stable over weeks in adults (42), we revisited the same regions 2 weeks later (Figure S2 D). Strikingly, we observed that a majority (67.8 ± 7.56%) of the same ECs retained their activity status (Figure 1 J-K; Movie 3) even though their dynamic properties were largely not retained (5.58 ± 6.51% retention of high frequency ECs and 0% retention of high average duration ECs) (Figure 1 L). Consistent with shorter term revisits, Ca^2+^ dynamics (frequency and average duration), though not conserved at the cellular level, were maintained over weeks at the network level (Figure 1 L-O). Lastly, the proportion of Ca^2+^ activity involving EC clusters was also conserved over weeks (Figure 1 P).

Collectively, our findings reveal that Ca^2+^ activity in the skin capillary plexus is dynamic during vascular homeostasis and spatially patterned by an active network of ECs with temporally conserved properties over time.

### Loss of endothelial Connexin 43 leads to chronically sustained Ca^2+^ activity and impairs long-term network regulation

Our findings that EC Ca^2+^ properties are conserved at the network level over time raise the question of how this is regulated molecularly. Gap junctions are intercellular channels that have established roles in EC communication and functional coupling (34, 49, 50). We hypothesized that maintenance of the active network is spatially coordinated by gap junctions. Connexin 43 is a gap junction protein both widely expressed in the vasculature, and the most expressed Connexin isoform in isolated skin ECs (Figure S3 A i) (51, 52). To investigate the role of Cx43 in the regulation of network Ca^2+^ activity, we combined our Ca^2+^ reporter with Cx43 knockout mice (Cx43^fl/fl^) to generate EC conditional knockouts (VECadCreER; Rosa26-CAG-LSL-GCaMP6s; LSL-H2B-mCherry; Cx43^fl/fl^ denoted as Cx43cKO) and compared them to littermate controls (VECadCreER; Rosa26-CAG-LSL-GCaMP6s; LSL-H2B-mCherry; Cx43^+/+^).

Cx43cKO and control mice were induced over 4 consecutive days (2 mg Tamoxifen daily) and then imaged 3 days later (1 week following the first Tamoxifen injection, referred to as Day 0). Surprisingly, we observed an increase in capillary plexus Ca^2+^ activity in Cx43cKO mice compared to controls (Figures 2 A-B; Movie 4). This included a significant increase in the proportion of active ECs (68.6 ± 3.71% in Cx43cKO mice vs. 50.8 ± 8.47% in control mice) (Figure 2 C). We also observed that the active network in Cx43cKO mice displayed significantly increased frequency (0.646 ± 0.586 events/min in the Cx43cKO mice vs. 0.473 ± 0.467 events/min in control mice) and a significant increase in the average duration of Ca^2+^ events per cell (62.5 ± 119 seconds in the Cx43cKO mice vs. 8.00 ± 7.33 seconds in control mice) (Figures 2 D-E). Furthermore, we found a group of ECs which displayed significantly long-lasting Ca^2+^ events ranging between 2 and 17 minutes, a behavior of persistent activity not detected in control mice (19.3 ± 6.51% of active ECs in Cx43cKO mice vs. 0.324 ± 0.305% of active ECs in control mice) (Figure 2 F). In contrast to control mice where EC clusters were active simultaneously only for a few frames, Cx43cKO showed “persistently active” EC clusters (61.7 ± 17.3% involved at least 2 cells), simultaneously active for the entire duration of recording (Figure S3 B-D). To understand whether changes in expression of other vascular Connexins (50) may influence Ca^2+^ activity in Cx43cKO mice, we used qPCR to analyze Connexins 37 and 40 expression in isolated skin ECs. We found their transcripts to be similarly expressed in Cx43cKO versus control samples, indicating an absence of compensatory changes in expression (Figure S3 A ii).

**Figure 2.**
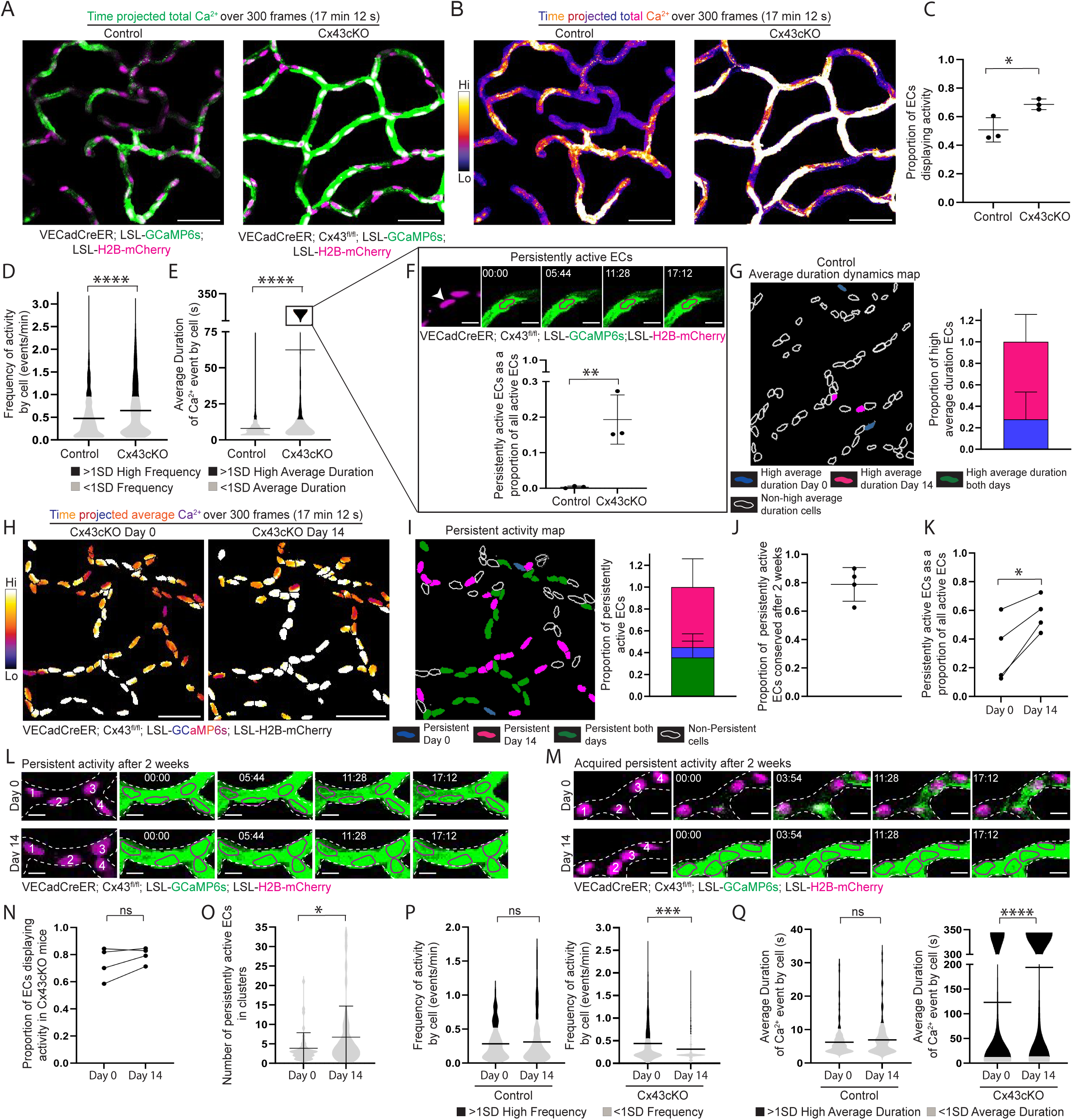
Loss of endothelial Connexin 43 leads to sustained Ca^2+^ dynamics and impairs long-term network regulation. **(A)** Max intensity projection of GCaMP6s signal (green) with H2B-mCherry signal (magenta) from 300 frame (17 minutes 12 seconds) recording of skin capillary ECs from control (VECadCreER; Rosa26-CAG-LSL-GCaMP6s; LSL-H2B-mCherry) and Cx43cKO (VECadCreER; Cx43^fl/fl^; Rosa26-CAG-LSL-GCaMP6s; LSL-H2B-mCherry) mice (scale bar: 50 µm). **(B)** Max intensity projection of GCaMP6s signal from (A) represented with fire lookup table. Fire lookup table allows for easier visualization of changes in Ca^2+^ signaling intensity. Color scale indicates GCaMP6s signal over total duration of recording (scale bar: 50 µm). **(C)** Proportion of active ECs in control and Cx43cKO mice. P=0.0288, unpaired t-test; *n=*10 regions from 3 mice each for control and Cx43cKO. **(D)** and **(E)** Frequency of Ca^2+^ activity (events/min) and average duration (s) of a Ca^2+^ event per cell respectively for control and Cx43cKO mice; black (>1SD high activity) and grey (<1SD activity). *n=*570 active ECs from 3 control mice and 837 active ECs from 3 Cx43cKO mice; P<0.0001 for both, unpaired t-test. **(F)** *Top*: Representative EC (drawn in magenta outline corresponding with H2B-mCherry label) displaying persistent activity in Cx43cKO mouse (scale bar: 10 µm). *Bottom*: Persistently active ECs as a proportion of total active ECs in control and Cx43cKO mice; P=0.009, unpaired t-test. **(G)** *Left:* Representative average duration cells map of control ECs revisited after 14 days. High average duration ECs on Day 0 (blue), Day 14 (magenta) or both days (green), and non-high average duration ECs (white outline) *Right:* Bar graph of high average duration ECs on day 0, 14, or both days. *n=*4 regions from 3 mice **(H)** Average intensity projections of GCaMP6s signal over H2B-mCherry region for single-cell resolution (fire lookup table) in Cx43cKO ECs revisited after 14 days (scale bar: 50 µm). Color scale indicates average GCaMP6s signal over total duration of recording. Average intensity projection provides visualization of persistently active ECs in white. **(I)** *Left*: Persistent activity map of ECs revisited after 14 days. Persistently active ECs on Day 0 (blue), Day 14 (magenta) or both days (green), and non-persistently active ECs (white outline). *Right*: Bar graph of persistently active ECs on Day 0, 14, or both days. *n=7* regions from 4 mice. **(J)** Proportion of persistently active ECs conserved after 14 days among revisited mice. **(K)** Persistently active ECs as a proportion of all active ECs, in mice revisited after 14 days; P=0.023, paired t-test. *n=7* regions from 4 mice. **(L)** Representative image of ECs (numbered and drawn in magenta outline of H2B-mCherry signal) maintaining persistent activity after 14 days in Cx43cKO mouse (scale bar: 10 µm). **(M)** Representative image of ECs gaining persistent activity on Day 14 (scale bar: 10 µm). **(N)** Proportion of active ECs from Cx43cKO mice on Day 0 and Day 14; ns: P>0.05, t-test, *n=*532 total cells from 4 mice. **(O)** Number of persistently active ECs in clusters, for regions revisited after 14 days; P=0.047, unpaired t-test. **(P)** Frequency of Ca^2+^ activity (events/min) and average duration **(Q)** of Ca^2+^ events per cell (s) on Day 0 and Day 14 of revisited ECs in control and Cx43cKO mice; black (>1SD high activity) and grey (<1SD activity). P<0.0001, unpaired t-test in Cx43cKO mice; *n=*399 active cells from 4 mice. ns= P>0.05, unpaired t-test in control mice; *n=*126 active cells from 3 mice.

We next asked whether spatiotemporal Ca^2+^ activity and network-level dynamics of ECs are conserved over time in Cx43cKO mice, as was observed under physiological conditions. Revisiting the same cells in the same Cx43cKO mice, we found that the activity status of ECs was spatially conserved on a single cell level after 14 days (78.4 ± 4.84% of ECs retained their status) (Figure S3 E and F). Furthermore, persistent Ca^2+^ activity was also conserved on a cellular level after 14 days (78.9 ± 11.5% of persistently active ECs on Day 0 retained their behavior on Day 14) (Figures 2 G-J and L; Movie 5). While there was no change in the proportion of active ECs after 14 days (Figure 2 N), we observed an increase in the proportion of persistently active cells (32.2 ± 22.9% on Day 0 vs. 57.3 ± 12.2% of active ECs on Day 14) (Figures 2 K and M; Movie 5). This was accompanied by an increase in the cluster size of ECs with persistent activity (from 3.85 ± 4 cells on Day 0 to 6.73 ± 7.99 cells on Day 14) (Figure 2 O). While control mice demonstrated conserved dynamics across the network after 14 days, Cx43cKO mice showed a significant increase in average duration (123 ± 156 seconds on Day 0 vs. 194 ± 165 seconds on Day 14) and a significant decrease in frequency (0.445 ± 0.432 events/min on Day 0 vs. 0.319 ± 0.322 events/min on Day 14) per cell, consistent with the observed increase in the number of persistently active cells (Figures 2 P and Q).

Our findings show that loss of Cx43 increases plexus-wide Ca^2+^ activity, enriching for a population of persistently active ECs. In addition, we found that loss of Cx43 maintains spatial patterning over weeks, but biases ECs towards longer-lasting Ca^2+^ activity, and accordingly, persistently active ECs become a larger fraction of the vascular plexus over time.

### L-type Voltage Gated Ca^2+^ channels (VGCCs) sustain elevated EC Ca^2+^ dynamics following loss of endothelial Connexin 43

Upon loss of Cx43, we captured an unexpected increase in Ca^2+^ activity across the capillary EC network. To identify what molecular mediators are responsible for this signaling phenotype, we tested a small panel of well-known inhibitors of ion channels involved in Ca^2+^ entry. Specifically, we topically applied GSK219 (TRPV4 inhibitor) (17), Mibefradil (T-type VGCC inhibitor) (53), Nifedipine (L-type VGCC inhibitor) (54), Senicapoc (Gardos channel inhibitor) (55), and DMSO vehicle in Cx43cKO mice, and imaged the same capillary regions before and after treatment (within 30-60 minutes) (17, 53, 56). Interestingly, we observed that nifedipine appeared to largely decrease total and persistent Ca^2+^ activity in Cx43cKO mice relative to the other compounds (Figure S4). In addition to L-type VGCCs, T-type VGCCs (54) and KCa3.1 channels (55) are possible other targets of nifedipine. Given their inhibition by Mibefradil or Senicapoc did not have any effect on Ca^2+^ activity in Cx43cKO mice, L-type VGCCs specifically emerge as molecular mediators of increased Ca^2+^ activity in ECs after loss of Cx43. To further test for specificity of L-type VGCC inhibition in driving the EC Ca^2+^ reduction after loss of Cx43, we also used a different L-type VGCC inhibitor, Verapamil, and lower doses of nifedipine (10 and 100 times reduced in concentration), all of which decreased Ca^2+^ activity in Cx43cKO mice (Figure S5 A).

Nifedipine is an inhibitor of L-type VGCCs (54, 56, 58, 59). To first test whether nifedipine would alter capillary Ca^2+^ activity during homeostasis, we used control (VECadCreER; Rosa26-CAG-LSL-GCaMP6s; LSL-H2B-mCherry) mice and compared the proportion of active ECs, as well as mean values of frequency and average duration by cell per mouse, before and after treatment with nifedipine or DMSO. We did not observe a significant difference in proportion of active ECs (13.7% ± 18.2% decrease after nifedipine treatment vs. 20.2% ± 15.0% decrease after DMSO treatment), frequency (20.6% ± 20.5% decrease after nifedipine treatment vs. 17.8% ± 17.9% decrease after DMSO treatment), or average duration (3.08% ± 8.73% decrease after nifedipine treatment vs. 11.8% ± 7.47 % decrease after DMSO treatment) (Figures 3 A-C and E; Movie 6). In addition, distributions of Ca^2+^ event frequency and average duration per cell were also not different before or after nifedipine treatment, suggesting that L-type VGCC inhibition does not affect network-wide Ca^2+^ activity in control mice (Figures 3 D and F).

**Figure 3.**
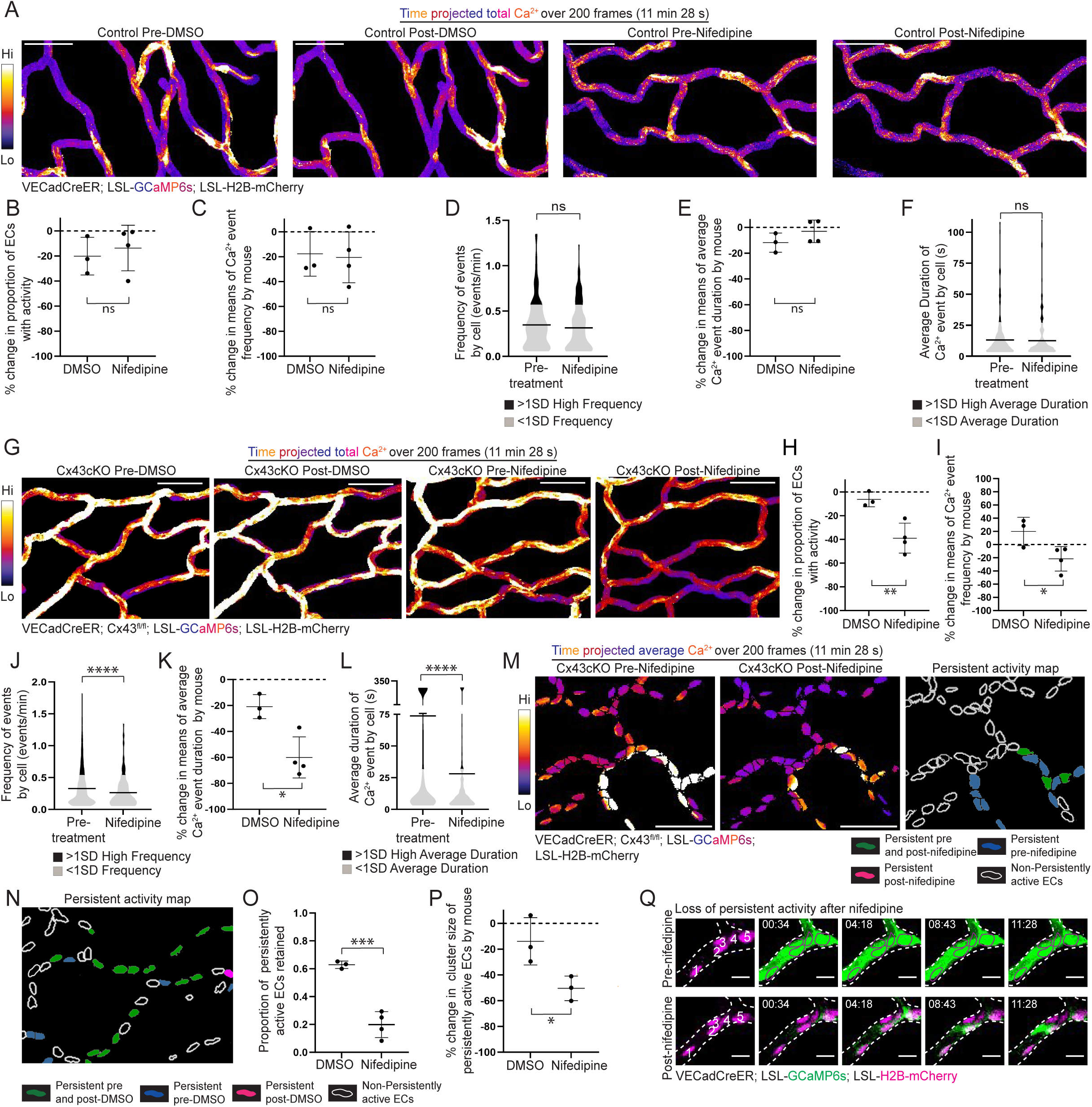
L-type VGCCs regulate sustained Ca2^+^ dynamics after Cx43cKO, but do not affect Ca^2+^ activity under physiological conditions. **(A)** Max intensity projection of GCaMP6s signal in control mice before and after treatment with either DMSO or nifedipine. GCaMP6s signal is represented with fire lookup table from 200 frame (11 minutes 28 seconds) recording of skin capillary ECs before and after treatment. Fire lookup table allows for easier visualization of changes in Ca^2+^ signaling intensity. Color scale indicates GCaMP6s signal over recording time (scale bar: 50 µm). **(B)** Percent change in proportion of active ECs in control mice revisited after treatment with either DMSO or nifedipine. ns=P>0.05, unpaired t-test; *n=* 5 regions from 3 mice for DMSO treatment and 6 regions from 4 mice for nifedipine treatment. **(C)** and **(E)** Percent change in frequency (events/min) and average duration (s) dynamics of ECs in control mice revisited after treatment with either DMSO or nifedipine (shown as percentage change of means, refer to Methods under Statistics and Reproducibility). ns=P>0.05, unpaired t-test. **(D)** and **(F)** Frequency of Ca^2+^ activity (events/min) and average duration (s) of Ca^2+^ events per cell of revisited control mice before and after nifedipine treatment. Black (>1SD high activity) and grey (<1SD activity); ns=P>0.05, unpaired t-test; *n=*187 active ECs pre-treatment and *n=*173 active ECs after treatment, from 4 mice**. (G)** Max intensity projection of GCaMP6s signal in Cx43cKO mice before and after treatment with either DMSO or nifedipine. (scale bar: 50 µm). **(H)** Percent change in proportion of active ECs in Cx43cKO mice revisited after treatment with either DMSO or nifedipine. P=0.0097, unpaired t-test; *n=*5 regions from 3 mice for DMSO treatment and 6 regions from 4 mice for nifedipine treatment. **(I)** and **(K)** Percent change in frequency (events/min) and average duration (s) dynamics of ECs in Cx43cKO mice revisited after treatment with either DMSO or nifedipine. P=0.04, unpaired t-test, for frequency plot. P=0.013, unpaired t-test, for average duration plot. **(J)** and **(L)** Frequency of Ca^2+^ activity (events/min) and average duration (s) of Ca^2+^ events per cell of revisited regions before and after nifedipine treatment. Black (>1SD high activity) and grey (<1SD activity); P<0.0001, unpaired t-test; *n=*358 active ECs pre-treatment and *n=*218 active ECs after treatment, from 4 mice. **(M)** *Left:* Average intensity projections of GCaMP6s signal (fire lookup table) over H2B-mCherry region for single-cell resolution, from regions revisited after nifedipine treatment (scale bar: 50 µm). Color scale indicates average GCaMP6s signal over total recording time. Average intensity projection improves visualization of persistently active ECs in white. *Right*: Map of persistently active ECs after nifedipine treatment. Persistently active ECs pre- and post-nifedipine (green), pre-nifedipine (blue), post-nifedipine (magenta) or non-persistently active ECs (white outline). **(N)** and **(O)** *Left*: Representative map of persistently active ECs in Cx43cKO mice revisited after treatment with DMSO. *Right*: Proportion of persistently active ECs retained after DMSO versus nifedipine treatments. *n=*5 regions from 3 mice for DMSO and 6 regions from 4 mice for nifedipine treatment. **(P)** Percent change in cluster size of persistently active ECs revisited after treatment with either DMSO or nifedipine; P=0.038, unpaired t-test. **(Q)** Representative images of ECs without persistent activity after nifedipine treatment in a Cx43cKO mouse (scale bar: 10 µm).

Next, to probe deeper into the effect of nifedipine treatment on Ca^2+^ activity in mice lacking EC Cx43, we repeated the experiments utilizing Cx43cKO mice with the H2B-mCherry nuclear reporter. Compared to DMSO treatment, nifedipine treatment led to decreased proportion of active ECs (39.0% ± 12.6% decrease after nifedipine treatment vs. 6.29% ± 6.23% decrease after DMSO treatment), frequency (21.5% ± 18.5% decrease after nifedipine treatment vs. a 20.0% ± 21.5% increase after DMSO treatment), and average duration (72.8% ± 37.0% decrease after nifedipine treatment vs. 29.3% ± 11% decrease after DMSO treatment) (Figures 3 G-I and K; Movie 7). In addition, the distributions of Ca^2+^ event frequency and average duration per cell were decreased after nifedipine treatment compared to pre-treatment (Figures 3 J and L). We next analyzed the effects of nifedipine on persistently active cells in Cx43cKO mice and confirmed a decrease in activity after treatment (19.9% ± 9.3% of persistently active cells retained after nifedipine treatment vs. 62.9% ± 2.65% after DMSO treatment) alongside a decrease in mean cluster size of persistently active ECs (50.5% ± 9.53% decrease after nifedipine treatment vs. 18.4% ± 14.0% decrease after DMSO treatment) (Figures 3 M-Q).

L-type VGCCs are known to be expressed in mural cells, including pericytes (60–63), but not endothelial cells (51, 52). To confirm this, we isolated skin ECs from Cx43cKO and control mice, extracted mRNA, and showed that neither the L-type VGCC main domain *CACNA1C* or its channel subunit *CACNA2D1* are expressed in ECs, while their expression was verified on isolated pericyte mRNA (Figure S5 B and C). All together, these data support a non-cell autonomous model of regulation whereby L-type VGCCs on neighboring cells mediate changes to a more persistent Ca^2+^ activity in capillary ECs after the loss of Cx43, but do not affect EC Ca^2+^ dynamics in control mice.

### Sustained Ca^2+^ activity after loss of endothelial Connexin 43 leads to vascular flow and barrier dysfunction

The elevated Ca^2+^ activity of Cx43cKO ECs over days to weeks was surprising since sustained cytosolic Ca^2+^ has been associated with cell death (64–66). This prompted us to assess EC number and endothelial architecture, but we found no change in cell density or vessel architecture even after revisiting the same cells in Cx43cKO mice after 14 days of sustained Ca^2+^ dynamics (Figure S6 A-C). Cx43 has also been shown to affect vascular flow and barrier function (67–75). We first investigated flow rate by injecting 150 kDa tetramethylrhodamine (TRITC) dextran into the vasculature of Cx43cKO and control mice and used line scanning as a method to track the number of cells passing through a region of the vessel over time (cell flux per second) while simultaneously recording Ca^2+^ activity (Figure 4 A). We tracked each vessel for a period of 5 minutes, at 2 second intervals, collecting both flow rate as well as Ca^2+^ signaling events and their durations (Figure 4 B). We observed a significant increase in average flow rate per vessel in Cx43cKO mice when compared to controls (42.5 ± 18 cells per second in Cx43cKO mice versus 28.9 ± 15.7 cells per second in control mice) (Figure 4 D and E; Figure S6 E). Intriguingly, we observed no apparent correlation between individual Ca^2+^ events and changes in local flow in the control mice, and no significant differences between changes in flow rate during signaling and non-signaling periods while tracking skin capillaries in homeostasis (Figure 4 B and C; Figure S7).

**Figure 4.**
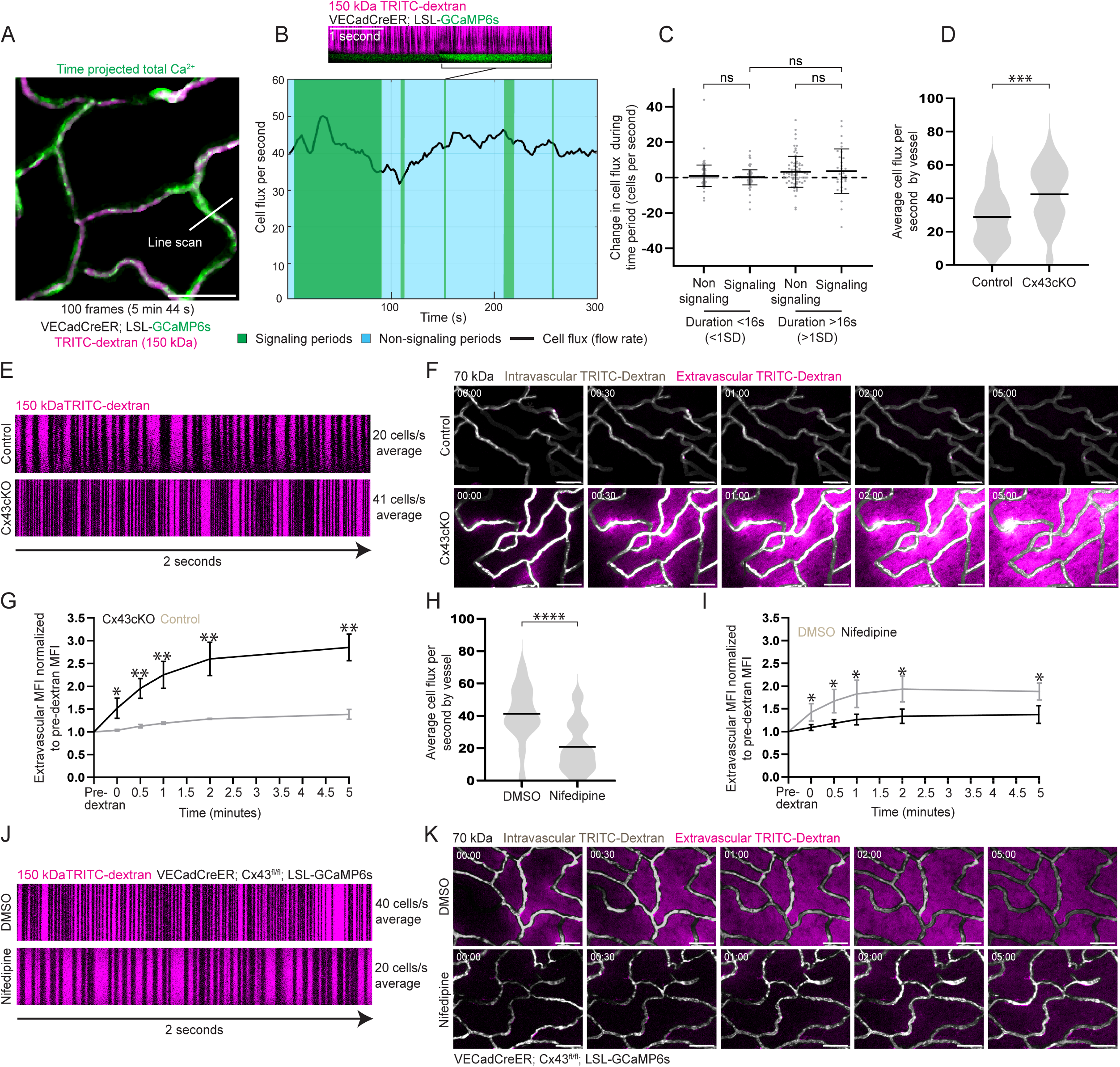
Sustained Ca^2+^ dynamics after Cx43cKO lead to vascular flow and barrier dysfunction. **(A)** Max intensity projection of GCaMP6s signal (green) with 150 kDa TRITC dextran (magenta) from 100 frame (5 minutes 44 seconds) recording, with the white line representing a line scanning region (scale bar: 50 µm). **(B)** *Top:* Representative line scan of a vessel over a 4 second period with Ca^2+^ events (green) and 150 kDa TRITC dextran for flow. Black lines in dextran represent individual cells passing through the line scan (scale bar: 1 s). *Bottom:* Graph representing cell flux per second or flow rate during line scanning of an individual vessel over a 5 min period. Flow rate (black line) is represented over signaling (green) and non-signaling (light blue) time periods. **(C)** Change in flow rate during non-signaling and signaling time periods (< and > 1SD in duration for each category). ns: P>0.05, unpaired t-test, *n=*45 vessels and 148 signaling events from 3 mice. **(D)** Average cell flux per second by vessel in control and Cx43cKO mice. P<0.001, unpaired t-test, *n=*45 vessels from 3 mice for each group. **(E)** Representative line scans of control and Cx43cKO vessel over a 2 second period. **(F)** Representative single time point images of 70 kDa intravascular dextran (grey) and extravascular dextran (magenta) (scale bar: 50 µm). **(G)** Line plots of extravascular MFI in control and Cx43cKO mice, normalized to pre-dextran MFI. P=0.020 at 0 min, P=0.003 at 0.5, 1, and 2 mins, and P=0.001 at 5 mins, unpaired t-tests; *n=* 3 mice each for control and Cx43cKO. **(H)** Average cell flux per second by vessel in Cx43cKO mice treated with DMSO or nifedipine. ns: P<0.0001, unpaired t-test, *n=*45 vessels from 3 mice for each group. **(I)** Line plots of extravascular MFI in DMSO and nifedipine treated Cx43cKO mice, normalized to pre-dextran MFI. P=0.02 at 0 mins, P=0.014 at 0.5 mins, P=0.017 at 1 min, P=0.015 at 2 mins, and P=0.018 at 5 mins, unpaired t-tests; *n=* 4 mice with nifedipine treatment and 3 control mice. **(J)** Representative line scans of Cx43cKO mouse after treatment with DMSO or nifedipine. **(K)** Representative single time point images of 70 kDa intravascular dextran (grey) and extravascular dextran (magenta) after treatment with DMSO or nifedipine (scale bar: 50 µm).

To test the potential impact of Cx43cKO on vessel permeability, we utilized a small size tetramethylrhodamine (TRITC)-dextran (70 kDa), known to be impermeable in skin capillaries during homeostasis (76, 77). We discovered that within seconds of injection, 70 kDa dextran was found in the interstitial space of skin capillaries of Cx43cKO mice but not in control mice (Figure 4 F and G). This is in contrast to the larger size dextran (150 kDa) used to assess flow, which did not leak into the interstitial space in Cx43cKO mice (Figure S6 D), indicating Cx43cKO’s barrier phenotype is size-specific and does not reflect an entirely dysfunctional endothelial barrier.

These defects in flow and barrier function prompted the question as to whether they are downstream of elevated Ca^2+^ signaling driven by loss of Cx43. Therefore, we employed nifedipine treatment, since we showed it rescued the Ca^2+^ elevation in Cx43cKO mice. Excitingly, upon nifedipine treatment, the dysregulated vascular flow was reverted back to physiological patterns (41.2 ± 17.1 cells per second in Cx43cKO mice treated with DMSO versus 20.8 ± 15.3 cells per second in Cx43cKO mice treated with nifedipine) (Figure 4 H and J; Figure S6 F). Leakage of 70 kDa dextran in the interstitial space of the skin was also significantly reduced with nifedipine treatment, even though we did observe non-specific reduction of permeability with DMSO treatment (Figure 4 I and K).

Altogether, our results demonstrate that sustained Ca^2+^ signaling after loss of EC Cx43 drives vascular flow dysregulation and barrier dysfunction (Figure 5).

**Figure 5.**
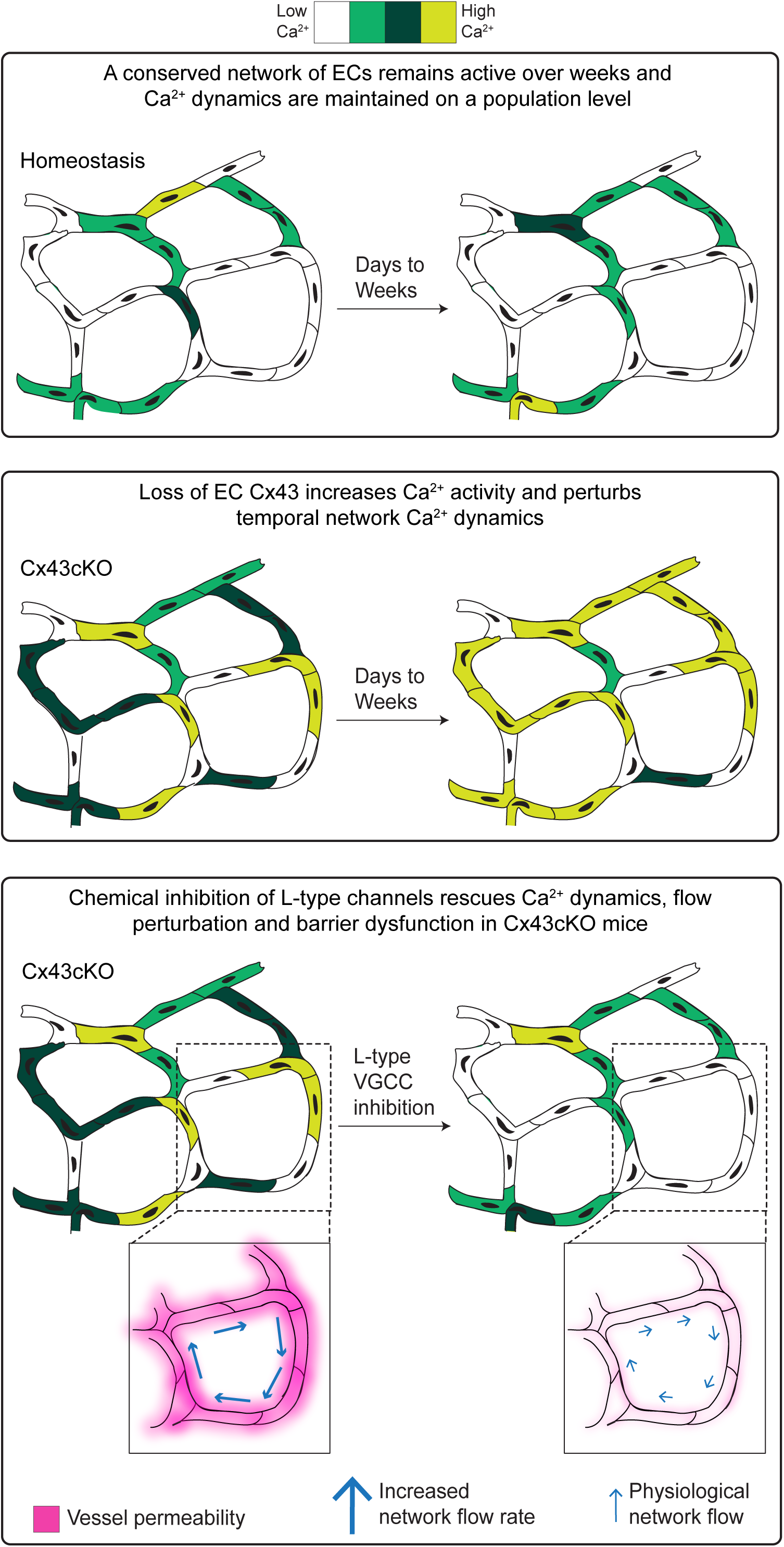
A conserved network orchestrates Ca^2+^ patterning in capillary ECs over days to weeks and is spatiotemporally and functionally regulated by Cx43. Model depicting the spatiotemporal characteristics of Ca^2+^ signaling organization in capillary ECs. A spatiotemporally conserved network of ECs orchestrates tissue-wide Ca^2+^ activity over days to weeks and maintains Ca^2+^ dynamics at the population level. Loss of Cx43 increases Ca^2+^ activity and impairs long-term temporal regulation of the EC network, converging on a persistently active phenotype. Sustained Ca^2+^ activity leads to flow and barrier dysfunction. L-type VGCC inhibition non cell-autonomously restores Ca^2+^ activity, flow, and barrier function in Cx43cKO mice to physiological patterns.

## Discussion

How Ca^2+^ activity is organized, maintained, and regulated across a vascular plexus in its native, unperturbed tissue environment is understudied. To our knowledge, our study represents the first time that Ca^2+^ activity of the same *in vivo* populations of mammalian ECs has been longitudinally tracked over minutes to days to weeks, demonstrating unexpected spatiotemporal patterns of endothelial Ca^2+^ activity and dynamics. Our tissue-wide analysis reveals that a network of active cells, conserved over time, orchestrates network Ca^2+^ activity (Figure 5). Coordinated Ca^2+^ activity as a baseline in adult capillary homeostasis highlights that organized cell networks control Ca^2+^ activity at the tissue-level, and reinforces the notion of the endothelium being a functional syncytium (29, 34, 78). Furthermore, we discovered that Cx43 functions to constrain endothelial Ca^2+^ activity within a dynamic range and maintain network signaling dynamics over the long-term (Figure 5). Our findings elucidate a role for Cx43 as a mediator of cell-cell communication on larger spatial and temporal scales than previously reported. Furthermore, we found that non-cell autonomous inhibition of L-type VGCCs was able to rescue EC Ca^2+^ signaling patterns, barrier function, and blood flow, revealing that a dynamic range of Ca^2+^ signaling across the EC network allows maintenance of physiological functions (Figure 5).

In this study, we discovered and quantified the organization and regulation of homeostatic Ca^2+^ activity in skin vasculature through our unique ability to resolve tissue-wide Ca^2+^ dynamics with single cell resolution *in an intact, uninjured mouse*. The plexus-wide Ca^2+^ activity in skin endothelial cells in physiological conditions (Figure 1) is reminiscent of Ca^2+^ activity observed *in vivo* in other tissues, including mouse skin epidermal stem cells and brain capillaries (4, 17). However, our uncovering of the spatiotemporal conservation of Ca^2+^ activity and its large-scale coordination emerges as critical features of skin capillaries (Figure 1). Specifically, the discovery that EC network activity status is maintained on a cellular level and that the distribution of Ca^2+^ dynamics is conserved on a population level are novel findings across organs and organisms to date. As such, a conceptual advance of our study is our demonstration that Ca^2+^ signaling in capillary endothelial cells is maintained at the network level over time. While large-scale coordination between ECs has been shown in periods of extensive vascular remodeling, such as development and injury (40, 42, 43, 79, 80), our work establishes an unexpected paradigm that Ca^2+^ activity in capillaries is both orchestrated at a tissue-wide scale and is non-random in its spatial patterning. Heterogeneity of EC Ca^2+^ activity, which is conserved at a single-cell level over days to weeks, also suggests cellular heterogeneity in the same vascular compartment. This builds upon findings of EC heterogeneity in the same vascular compartment, across the vascular tree, and between organs, as a way for the endothelium to adapt to local environments (76, 81–88). Our findings may further indicate differences in the microenvironment and/or decision-making on a cell-to-cell basis in a capillary plexus.

We find that the gap junction protein Cx43 regulates the active network on a temporal scale by influencing network coordination and maintenance of Ca^2+^ dynamics over time (Figure 2). Gap junctions and connexins have been described as intercellular channels that allow for local cell-cell communication in all tissues (32, 33). However, after loss of Cx43 in ECs, there is a loss of temporal maintenance of population-level behaviors that leads to increasing proportion of persistent signaling ECs over weeks, emphasizing a novel long-range, tissue-level regulatory role of Cx43 in temporal maintenance of plexus-wide EC Ca^2+^ signaling.

Ca^2+^ activity in the Cx43cKO capillary plexus converges towards more persistent Ca^2+^ activity without any cell death, while maintaining similar proportions of active ECs (Figure S6). Given the role of elevated, sustained cytosolic Ca^2+^ in apoptosis and cell death (64–66), we anticipate that other mechanisms, including the protective effects of PMCA (Plasma Membrane Ca^2+^-ATPase) and SERCA (Sarco/Endoplasmic Reticulum Ca^2+^ ATPase) in reducing Ca^2+^ overload to promote cell survival, might function to keep high concentrations of intracellular Ca^2+^ in check (89, 90). The phenotype of persistently active skin capillary ECs in Cx43cKO mice (Figure 2) also contrasts with published *in vitro* work in ECs, where pharmacological pan-inhibition of gap junctions, or genetic knockout of different Connexin isoforms, decreases Ca^2+^ activity (26, 29, 35, 91–94). Our data may indicate a more complex regulation of EC Ca^2+^ signaling with the EC network structurally intact. Moreover, *ex vivo* approaches have shown that Ca^2+^ activity and spread recorded between adjacent alveolar capillaries is abolished in both EC-specific Cx43 knockout mice and mimetic peptide blocking of Cx43 function (69). We characterized Ca^2+^ signaling across the global vasculature in a native, unperturbed context, which may explain the differences in our findings compared to *ex vivo* models that more closely mimic injury contexts. Prior to this study, Ca^2+^ activity has not been clearly visualized *in vivo* after the loss of gap junctions. However, the reported vascular phenotypes in studies involving endothelial conditional deletion of Cx43, Cx37, and Cx40 include sustained vessel dilation and increase of plasma nitric oxide levels (34, 73). These phenotypes are associated with an increase in EC Ca^2+^ activity, and therefore we believe that the differences we observed in skin ECs compared to previously described systems upon loss of Cx43 reflect a combination of tissue-specificity, the physiological, native context and the unperturbed nature of the vascular architecture.

The interplay we observed between L-type VGCC activity and Cx43 dysregulation (Figure 3) provides insight into how heterotypic cell interactions can reshape the capillary Ca^2+^ landscape. We propose a model whereby the loss of Cx43 in capillary ECs allows for activation of L-type VGCCs on neighboring cells, which then non-cell autonomously regulate EC Ca^2+^ dynamics. Cx43 gap junctions have been implicated in maintaining and altering membrane potential through electrical coupling and ion exchange functions (95, 96). In this way, loss of Cx43 in ECs could influence membrane potential and L-type VGCC activation in neighboring cells, which then through ‘inside out’ signaling mechanisms or paracrine factors may influence EC Ca^2+^ signaling (31, 97, 98). Since pericytes are known to be coupled electrically with ECs through gap junctions, can influence EC Ca^2+^ activity in other tissues, and are widespread in skin (60–63, 99–102) (Figure S5 C), they are one possible candidate. However, we cannot rule out the involvement of other neighboring cell types including macrophages (103) and fibroblasts (104). Future work is needed to fully understand the role of voltage gated channels in regulating the skin capillary plexus.

Our discovery of sustained Ca^2+^ signaling after loss of Cx43 leading to flow and barrier dysfunction indicates that a set dynamic range of Ca^2+^ signaling across the network is necessary for homeostatic maintenance of vascular function, which when exceeded, leads to global flow dysregulation and barrier dysfunction. Ca^2+^ signaling in brain capillaries has been described to drive local blood flow changes (17, 23, 24). Our findings that homeostatic Ca^2+^ signaling is not correlated with apparent blood flow changes may indicate tissue-specific differences in Ca^2+^ regulation of flow and emphasize differences between central vasculature and regulation of general, peripheral vascular beds. In regard to vascular permeability, the direct role of elevated cytosolic Ca^2+^ in disrupting endothelial barrier integrity through RhoA-ROCK1 activation, VE-Cadherin junction disassembly, and cytoskeletal changes has been explored in past studies (105–109). We wonder whether similar mechanisms may drive changes in vascular permeability after sustained Ca^2+^ elevation in the skin vasculature. Increased shear stress can also lead to EC junctional changes (110, 111), and so the reduction in permeability after L-type VGCC inhibition in Cx43cKO mice may be directly influenced by the decrease in vascular flow rate.

In conclusion, our work provides unprecedented spatial and temporal intravital imaging at single cell resolution to define the spatiotemporally conserved networks of capillary EC Ca^2+^ signaling and reveals that vascular function is compromised when this signaling is altered.

### Limitations of the study

In this study, we investigated the spatiotemporal characteristics and regulation of tissue-level Ca^2+^ activity across capillary ECs *in vivo*, focusing on the skin. One consideration is using a pan-endothelial driver (VECadherin promoter) that does not allow us to discern EC heterogeneity or perturb specific subpopulations of ECs. Drug treatment absorption into the skin can also be variable. This issue was addressed by assessing multiple regions of the plexus within the same mice before and after treatments.

## Materials and Methods

### Mice

*VECadCreER* (112) mice were obtained from Ralf Adams (Max Planck, Muster, Germany), *Cx43^fl/fl^* (73, 113), *Rosa26-CAG-LSL-GCaMP6s* (114), *Rosa26-CAG-LSL-H2B-mCherry* (115), *aSMA-RFP* (116) and *LSL-tdTomato* (114) mice were obtained from The Jackson Laboratory. All experimental mice were bred to a mixed CD1 albino background. Mice from experimental and control groups were randomly selected from either sex for live imaging experiments, and were within an adult age range of 2–4 months. The experiments were not randomized, and all experimentation involving animals were performed under the approval of the Yale School of Medicine Institutional Animal Care and Use Committee (IACUC).

### In vivo imaging

Imaging procedures were adapted from those previously described (117). An isoflurane chamber was used to anesthetize mice, and then the mice were transferred to the imaging stage on a 37 °C heating pad. Mice were maintained on anesthesia throughout the course of recording with vaporized isoflurane delivered by a nose cone (1.25% in oxygen and air). The right paw was mounted on a custom-made stage and a glass coverslip was placed directly against the flat part of the paw. Image stacks were acquired with a LaVision TriM Scope II (LaVision Biotec) laser scanning microscope with a Chameleon Vision II (Coherent) two-photon laser (using 940 nm for live imaging) and a Chameleon Discovery (Coherent) two-photon laser (using 1120 nm for live imaging). A Nikon 25x/1 water immersion objective was used. Optical sections were scanned with a field of view of 0.3 x 0.3 mm^2^. For all timelapse movies, the live mouse remained anesthetized for the length of the experiment and serial optical sections (3 μm steps for a total stack of 12 μm) were captured at intervals of 3.44 seconds for a total of 200 (11 minutes 28 seconds for topical pharmacological experiments) or 300 frames (17 minutes 12 seconds total recording time).

For revisits, the same region of live mouse skin was imaged across intervals of multiple days, and anatomical features of the paw, such as mouse digits, were used as landmarks for finding the same location.

### Image analysis

Raw image stacks were imported into FIJI (ImageJ, National Institutes of Health) for analysis. Max or average (to select for persistently active behaviors specifically) Z stacks of sequential optical sections were generated. Translational motion correction of timelapse movies was performed in Imaris software.

Analysis of Ca^2+^ for individual ECs: Segmentation of the H2B-mCherry signal was performed via the threshold and masking functions on FIJI. Through a custom MATLAB pipeline, the GCaMP6s fluorescence intensity of each masked region of interest (ROI as a proxy for a cell) every 100 frames was normalized to the minimum fluorescence intensity over that period (Figure S2). Any change in fluorescence over a threshold (50% increase above the minimum fluorescence intensity) was treated as a Ca^2+^ event, and the duration of each event, as well as the number of events per ROI was calculated (Figure S2). To determine persistent signaling, a 50% change in minimum fluorescence intensity cannot be used because the mean MFI within the cell remains very high for the duration of imaging. In this case, any EC with an MFI exceeding 170 (out of a maximum of 255), was treated as signaling for the duration it exceeded that threshold. Frequency is described as the number of events per minute, with the lowest possible frequency by cell of 0.0588 events/minute (1 event over the 17 minutes 12 seconds recording duration) or 0.001 Hz. Average duration of Ca^2+^ events was analyzed per EC by dividing total signaling duration by the number of Ca^2+^ events per cell.

Analysis of Ca^2+^ for EC clusters: Segmentation of the GCaMP6s signal with manual masking was used to segment the vessel surface. Using our custom MATLAB pipeline, we divided the masked vessel surface into individual grids, 0.0076 x 0.0076 mm^2^ and the fluorescence intensity of each grid ROI was normalized every 100 frames by taking the minimum fluorescence value over each period (Figure S2). Any change in GCaMP6s fluorescence over 50% of the minimum fluorescence intensity was identified as a Ca^2+^ event, and the duration of each event, as well as the number of events per ROI was calculated. To analyze EC clusters, adjacent grid ROIs that displayed Ca^2+^ events either at the same frame or one frame apart were considered part of the cluster. The maximum number of grids with connected activity was represented, and the H2B-mCherry segmented ROIs were added to each set of connected grids to determine the number of ECs involved in each cluster (Figure S2).

Analysis and visualization of Ca^2+^ events by cell: A custom MATLAB pipeline was generated with assistance from ChatGPT-5 that takes an Excel spreadsheet of per-cell signaling event intervals, converts the event timing into seconds using the user-supplied frame time, and builds a binary cell × time activity raster showing where each cell is “on” during its event windows. The pipeline output was validated against cell signaling durations calculated manually to ensure the pipeline was functioning as anticipated. The pipeline then generates three visualizations labeled in seconds: (1) a raster heatmap of on/off signaling over time, (2) stacked per-cell “wave” traces, and (3) a bar plot of per-cell event frequency.

Analysis of vessel architecture: The capillary plexus in Cx43cKO mice was imaged with fine serial optical sections (0.5 μm steps for a total stack of 40 μm) on Day 0 and 14. The max projection of the optical sections was then used for subsequent analysis. A custom MATLAB script was generated with assistance from ChatGPT-5 to automatically identify fully enclosed vascular loops from a binary vessel mask. Each loop was labeled with a unique ID and its area, perimeter, centroid, and equivalent diameter were measured using MATLAB’s *regionprops* function. The pipeline was validated against regions where the parameters of loops were manually calculated.

Cell density: Endothelial cells were imaged with fine serial optical sections of 0.5 µm steps across 20-25 µm, on Day 0 and 14. Image stacks with uniform illumination across the scan area were chosen for analysis and were maximally projected for further segmentation. Nuclear masks were generated from the projected stacks using Cellpose-SAM (118) while the vascular skeleton was generated from a custom Python script. Each masked endothelial nuclear centroid was snapped onto the skeletonized vascular mask using a Euclidean distance transform and nuclei were retained only if the snapping distance was under 10 microns. Vessel segments were then detected as individual skeletonized paths between either branch points or endpoints using a connected neighborhood criterion and the length of each segment was calculated as a geodesic distance. For every segment, segment-wise linear endothelial cell density was then computed across the Day 0 and Day 14 revisits.

Line scan analysis: A custom MATLAB pipeline was generated with assistance from ChatGPT-5 that uses changes in MFI to understand cell flow (since cells are unlabeled in a TRITC dextran, black lines or MFI intensity valleys are cells moving through the dextran). The pipeline loads a RED 1-D intensity profile from a CSV (with user-selected columns if needed), prompts the user for peak/valley detection and block parameters, then identifies peaks and intensity-qualified valleys using prominence/distance/width rules. It converts the RED distance axis to a 0-based coordinate system, partitions the profile into fixed-width blocks (e.g., 600 pixels), and for each block counts valleys and computes valley-to-valley spacings. Next, it loads a GREEN MFI CSV, averages GREEN MFI within the same RED-defined blocks, and automatically labels each block as no event/before/during/after based on whether block-mean MFI exceeds a threshold (1.5× the minimum block mean). The script then generates an overall figure with blocks shaded by event label and valleys overlaid, optionally exports per-block PNGs, and writes an Excel workbook containing summary, distances, and event legend outputs. For each vessel, five random 2-second time blocks (out of 150 total) were manually checked with the pipeline output to ensure the user selected parameters accurately capture intensity valleys (cells).

Average flux was calculated as the average flow rate during an individual signaling or non-signaling time period. Vessel flux represents a smoothed trend of vessel flow rate per time point, averaged over each time point and its neighboring 4 time points (2 prior and 2 after). Change in flow rate during a time period was calculated by looking at the difference in flow rates between the first and last time points during that period.

### Quantitative PCR

RNA from isolated ECs was extracted using Qiagen RNeasy Plus Micro kit (74034). cDNA was made using SuperScript IV First-Strand Synthesis kit (Thermo Fisher 18091050). qPCR utilized FastStart Universal SYBR green Master (Sigma) on the CFX Connect Real-Time PCR Detection System (Bio-Rad). The set of primers used are as follows:

Connexin 37: forward, CCCACATCCGATACTGGGTG; reverse, CGAAGACGACCGTCCTCTG.

Connexin 40: forward, AGGGCTGAGCTTGCTTCTTA; reverse, TTAGTGCCAGTGTCGGGAAT.

Connexin 43: forward, GGTGATGAACAGTCTGCCTTTCG; reverse, GTGAGCCAAGTACAGGAGTGTG.

CACNA1C: forward, CGTTCTCATCCTGCTCAACACC; reverse, GAGCTTCAGGATCATCTCCACTG.

CACNA2D1: forward, GTGGAAGTGTGAGCGGATTGAC; reverse, TCGCTTGAACCAGGTGCTGGAA.

### Tamoxifen induction

To induce the expression of GCaMP6s, H2B-mCherry, and/or loss of Cx43 expression, *VECadCreER; Rosa26-CAG-LSL-GCaMP6s; LSL-H2B-mCherry* or *VECadCreER; Cx43^fl/fl^; Rosa26-CAG-LSL-GCaMP6s; LSL-H2B-mCherry* or *VECadCreER; Cx43^fl/fl^; Rosa26-CAG-LSL-GCaMP6s* or *VECadCreER; Rosa26-CAG-LSL-GCaMP6s* or *VeCadCreER; Cx43^fl/fl^* or *VeCadCreER; LSL-tdtomato* or *VeCadCreER* mice were given four doses of tamoxifen (2 mg in corn oil) 4, 5, 6, and 7 days before imaging or tissue collection by intraperitoneal injection (IP).

### Topical drug treatment

For each drug tested, 4-6 capillary regions were imaged for 200 frames (11 minutes 28 seconds). To inhibit T-type VGCCs, KCa3.1 channels, TRPV4, or L-type VGCCs, mibefradil, senicapoc, GSK219, verapamil, and nifedipine respectively were delivered topically to the paw skin. Nifedipine, verapamil, and GSK219 were dissolved in a 100 mg ml^−1^ stock solution in dimethyl sulfoxide (DMSO) and 30 μl of the mixture was spread evenly on the paw for 15 minutes. Mibefradil and senicapoc were dissolved in a 50 mg ml^−1^ stock solution in dimethyl sulfoxide (DMSO) and 50 μl of the mixture was spread evenly on the paw for 15 minutes. For lower doses of nifedipine treatment, the stock solution was diluted 10 or 100 times in DMSO before application. Regions in the paw were then revisited 30-60 minutes after application of the topical agents. A solution of 100% DMSO was used as vehicle control.

### Capillary Visualization in aSMA-RFP mice

For visualization of capillaries in aSMA-RFP mice, 15 mg/kg of 150 kDa fluorescein isothiocyanate (FITC) dextran was injected retro-orbitally, and a 0.28 x 0.28 mm^2^ region was imaged with a 40x water immersion objective, for a total depth of 50 microns with fine 0.5 µm steps. The max projection of 40 steps, or 20 µm of depth, was taken to capture the entire superficial capillary region.

### Vascular permeability

A 0.3 x 0.3 mm^2^ region was imaged for 100 frames (5 minutes 44 seconds) in *VECadCreER; Cx43^fl/fl^; Rosa26-CAG-LSL-GCaMP6s;* or *VECadCreER; Rosa26-CAG-LSL-GCaMP6s* anesthetized mice prior to dextran injection. 15 mg/kg of tetramethylrhodamine (TRITC) dextran (70 kDa or 150 kDa molecular weight) was injected retro-orbitally into each mouse. 10-15 seconds after retroorbital injection, the dextran flow was recorded in the same 0.3 x 0.3 mm^2^ region imaged before dextran injection, for a period of 100 frames using the Chameleon Discovery (Coherent) two-photon laser (1120 nm). For permeability rescue experiments, DMSO (vehicle) or nifedipine was spread evenly on the paw for 15 minutes, and then 15 mg/kg of TRITC-dextran (70 kDa molecular weight) was injected retro orbitally into each mouse.

For analysis of interstitial dextran leakage, segmentation of the TRITC-dextran with manual masking was used to segment the vascular structure. Pre-dextran, the GCaMP6S signal was used to segment the vessel surface. MFI of the regions outside the segmented vessel surface were calculated for each frame after dextran injection and normalized to the average extravascular MFI in the pre-dextran recording.

### Vascular flow and line scanning

A 0.3 x 0.3 mm^2^ region was imaged for 100 frames (5 minutes 44 seconds) in *VECadCreER; Cx43^fl/fl^; Rosa26-CAG-LSL-GCaMP6s;* or *VECadCreER; Rosa26-CAG-LSL-GCaMP6s* anesthetized mice prior to dextran injection. 15 mg/kg of TRITC-dextran (150 kDa molecular weight) was injected retro-orbitally into each mouse. Line scanning was conducted in the middle of individual vessels, at a frequency of 600 line scans over one optical section across each 2 second period. Each vessel was scanned for 150 time steps, or 300 seconds total. Vessels with large changes in baseline MFI during tracking were discarded for analysis, as this was an indication that the FOV shifted such that the vessel was not being captured throughout the imaging timeframe.

For flow rate rescue experiments, DMSO (vehicle) or nifedipine was spread evenly on the paw for 15 minutes, and then 15 mg/kg of TRITC-dextran (70 kDa molecular weight) was injected retro orbitally into each mouse prior to line scanning for 30-60 minutes after exposure.

### Endothelial cell isolation and sorting

*VECadCreER, VECadCreER; LSL-tdtomato,* and *VECadCreER; Cx43^fl/fl^* mice were induced with tamoxifen 1 week prior to harvesting tissue. Adult paws were collected and placed dermis side down in a 5 mg/ml dispase II solution (MilliporeSigma; 494207800) for 45 minutes at 37 °C. The epidermis was removed, and the dermis was minced finely and transferred to 0.25% collagenase IV (Sigma; C5138) in HBSS (GIBCO;14170-112) solution for at least 90 minutes at 37 °C. The tissue was then passed through an 18-gauge needle (BD; 305195) and washed in FACS Buffer (3% FBS, 2mM EDTA). The suspensions were then centrifuged at 350G for 10 minutes and filtered through a 40 µm filter (Falcon; 352340) before staining with anti-mouse CD31 APC (Biolegend;160210; 1/200) for sorting on a BD FACS AriaII. ECs were sorted based on expression of CD31 (and for *tdtomato* in *VECadCreER; LSL-tdtomato mice*). *aSMA-RFP* mice were used for harvesting tissue without any tamoxifen exposure, and the RFP was used to isolate cells.

### Statistics and reproducibility

GraphPad Prism software (GraphPad, Inc.) was used to perform statistical analyses (version 9.2). Parameters are reported in the figure legends, and comparisons between groups were made using paired or unpaired two-tailed Student’s *t* test (in short: t-test), or Chi-squared analysis for observed versus expected outcomes (df=1) in revisits. Normality was assumed in data distribution, but not formally tested. Differences between groups were considered significant at P < 0.05 and the data are presented as means ± SD.

High frequency or high average duration EC populations were designated as displaying activity 1 standard deviation (SD) greater than the mean of all active ECs. For control revisits, the value threshold for >1SD dynamics cells was calculated based on frequency and average duration distributions on Day 0. For Cx43cKO mice, the value threshold for >1SD dynamics cells was based on the frequency and average duration distributions of the control mice.

For Chi-squared analysis, expected values of ECs active on Day 0 and then also when revisited (‘random activity’ as the null hypothesis) were assigned based on the percentage of ECs displaying activity on Day 0 (58% for 24-hour revisits, 54% for 14-day revisits, and 74% for Cx43cKO 14-day revisits). These proportions were then compared to the percentage of cells that were observed to maintain their activity status upon revisiting.

To assess changes in frequency and average duration dynamics in revisited regions after pharmacological application of DMSO or nifedipine, the mean values of frequency and average duration were calculated for each mouse before and after treatment, and the percentage change between the pre-treatment and post-treatment values was compared between the two conditions.

## Acknowledgements

We thank all members of the Greco laboratory, especially S. Du and T. Xin, as well as B. Ehrlich, D. Greif, and Z. Sun (Yale), for critical feedback on the project as well as advice on tool development. This work was supported by National Institutes of Health grant nos. R01AR063663-12, R01AR072668-06, a Leo Foundation Grant LF-OC-23-001463, the Howard Hughes Medical Institute, and the European Research Council project no. 101167365 (all to VG), and a K99 award grant no. 5K99AR079555-02 (to CYK).

## Author Contributions

AS, CYK, JJM, and VG conceived the project and designed experiments. AS, VG, JJM, CYK and DMA wrote the manuscript. AS performed and/or participated in all experiments and analyzed the data. DGG and JLM helped build the analysis pipeline. CMM performed flow cytometry and qPCR experiments. DS and FX performed staining analyses. ZL conducted expression analyses from existing skin scRNA-seq databases. UR developed the pipeline for analyzing cell count and architecture. DGG, CMM, DS, FX, UR, ZL, and JLM provided critical feedback to the manuscript text and schematics.

---

**Figure S1.**
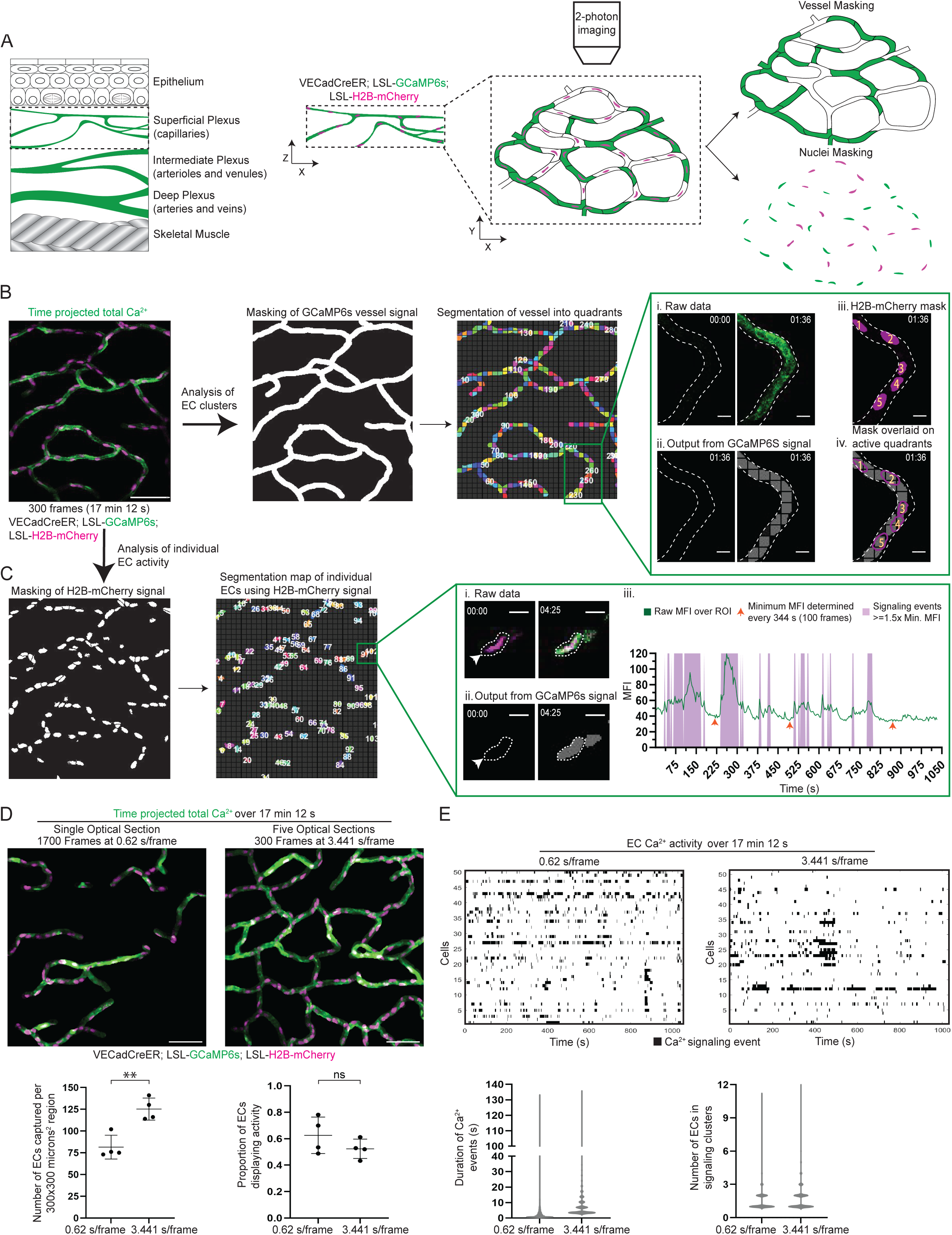
Schematic of intravital imaging and analysis platforms in Ca^2+^ sensor mice and analysis across kinetic scales. **(A)** *Left*: Cartoon representing skin in XZ plane, outlining the superficial capillary plexus. *Middle*: Cartoon of 2-photon imaging over the XY plane of the capillary region, with EC-specific GCaMP6s signal (green) and H2B-mCherry signal (magenta). *Right*: Representation of vessel and nuclei masking from imaged regions. **(B)** *Left:* Max intensity projection of GCaMP6s signal (green) with H2B-mCherry signal (magenta) from 300 frame (17 minutes 12 seconds) recording of skin capillary ECs (scale bar: 50 µm). *Middle:* Masking of vessels based on GCaMP6s signal, and segmentation of vessel mask into 0.0076 x 0.0076 mm^2^ quadrants. *Right:* **(i)** Representative images of a Ca^2+^ event, with the **(ii)** pipeline output displaying the number of quadrants involved (scale bar: 10 µm). An event is defined as at least a 50% increase above mean fluorescence intensity (MFI) over the minimum fluorescence for each quadrant. **(iii)** and **(iv)** Adding the H2B-mCherry signal over the pipeline output allows determining the number of ECs involved. **(C)** *Left:* Masking of nuclei based on H2B-mCherry signal as a proxy for individual ECs *Middle:* Segmentation of nuclei mask into individual regions of interest (ROIs) *Right:* **(i)** Representative images of a Ca^2+^ event (scale bar: 10 µm), with the **(ii)** pipeline output displaying an event. An event is defined as at least a 50% increase above MFI over the minimum fluorescence for each ROI. **(iii)** MFI (green) over the ROI over 300 frames of recording (17 minutes 12 seconds), with minimum MFI calculated every 100 frames (orange arrowheads), and overlaid output Ca^2+^ events (purple). **(D)** *Top:* Max intensity projection over 17 min 12 s recording at two frame rates: 0.62 s/frame over a single optical section or 3.44 s/frame over 5 optical sections (scale bar: 50 µm). *Bottom Left:* Number of ECs captured per 0.3 x 0.3 mm^2^ region when imaged at 0.62 s/frame and 3.44 s/frame. P=0.003, unpaired t-test; *n=*11 regions from 4 mice imaged at 0.62 s/frame and 14 regions from 4 mice imaged at 3.44 s/frame. *Bottom Right:* Proportion of active ECs in mice imaged at 0.62 s/frame and 3.44 s/frame. P=0.239, unpaired t-test; *n=*11 regions from 4 mice imaged at 0.62 s/frame and 14 regions from 4 mice imaged at 3.44 s/frame. **(E)** *Top:* Representative plots of Ca^2+^ signaling events (black) for 50 active ECs imaged at 0.62 seconds/frame and 50 active ECs imaged at 3.44 s/frame. *Bottom:* Duration of Ca^2+^ events and Number of ECs in signaling clusters in mice imaged at 0.62 s/frame and 3.44 s/frame; *n=*8,733 events from 4 mice imaged at 0.62 s/frame and 6,936 events from 4 mice imaged at 3.44 s/frame.

**Figure S2.**
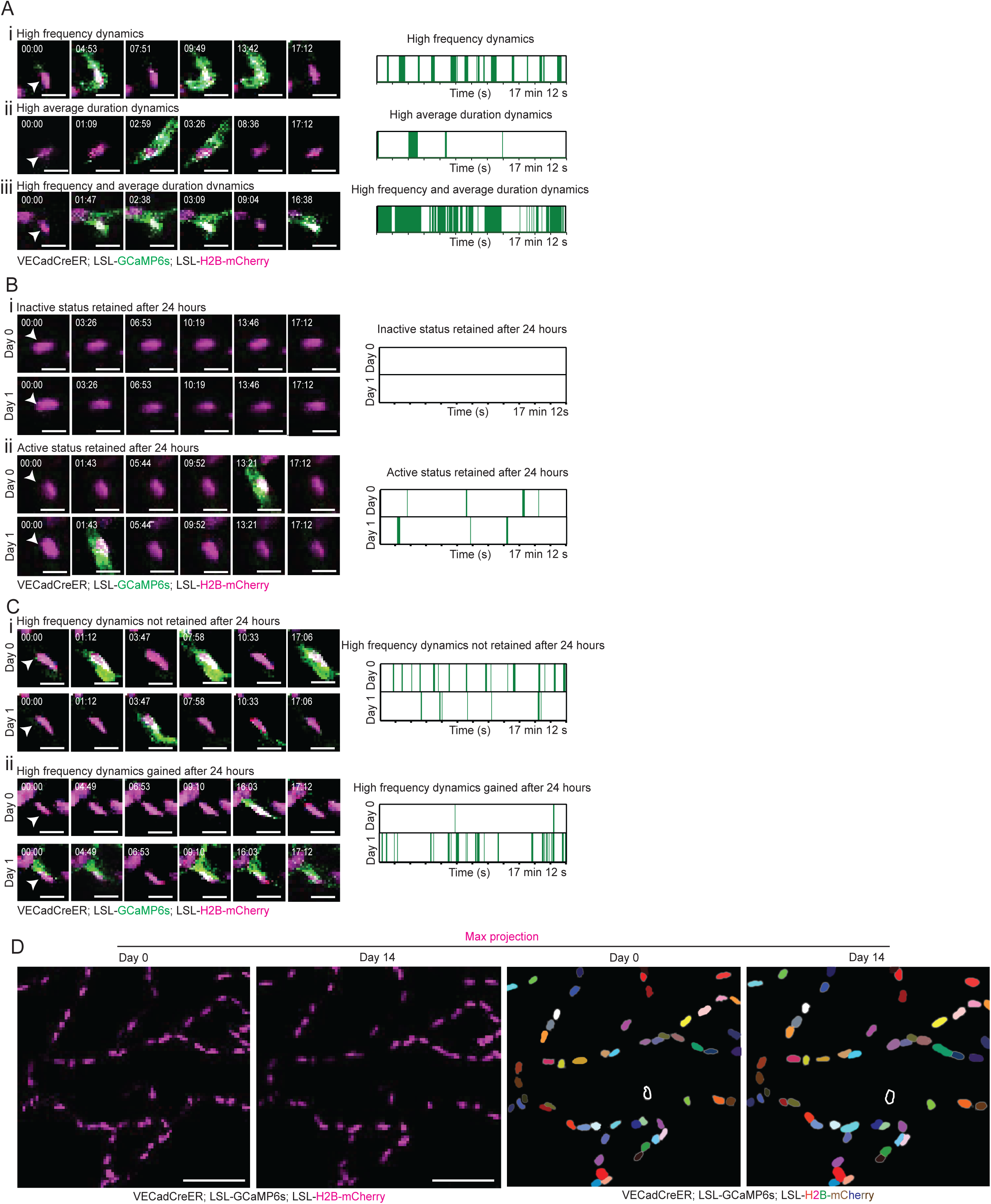
Ca2+ signaling dynamics and EC positional stability over time (A) **(i)** Representative image of EC (nuclei in magenta) displaying high frequency dynamics with plot showcasing Ca^2+^ events (green) and their durations over 300 frames (17 minutes 12 seconds) of recording (scale bar: 10 µm). **(ii)** EC displaying high average duration dynamics. **(iii)** EC displaying high frequency and high average duration dynamics. **(B) (i)** Representative image of EC displaying inactive status on Day 0 and again when revisited 24 hours later, (scale bar: 10 µm). **(ii)** EC displaying activity on Day 0 and again when revisited 24 hours later. **(C) (i)** Representative image of EC losing high frequency dynamics when revisited 24 hours later. (scale bar: 10 µm). **(ii)** EC gaining high frequency dynamics when revisited 24 hours later. (**D)** *Left:* Representative image of region revisited on Day 0 and Day 14 (scale bar: 50 µm). *Right:* ECs multi-color coded, with the same colors on Day 0 and Day 14 corresponding to the same EC.

**Figure S3.**
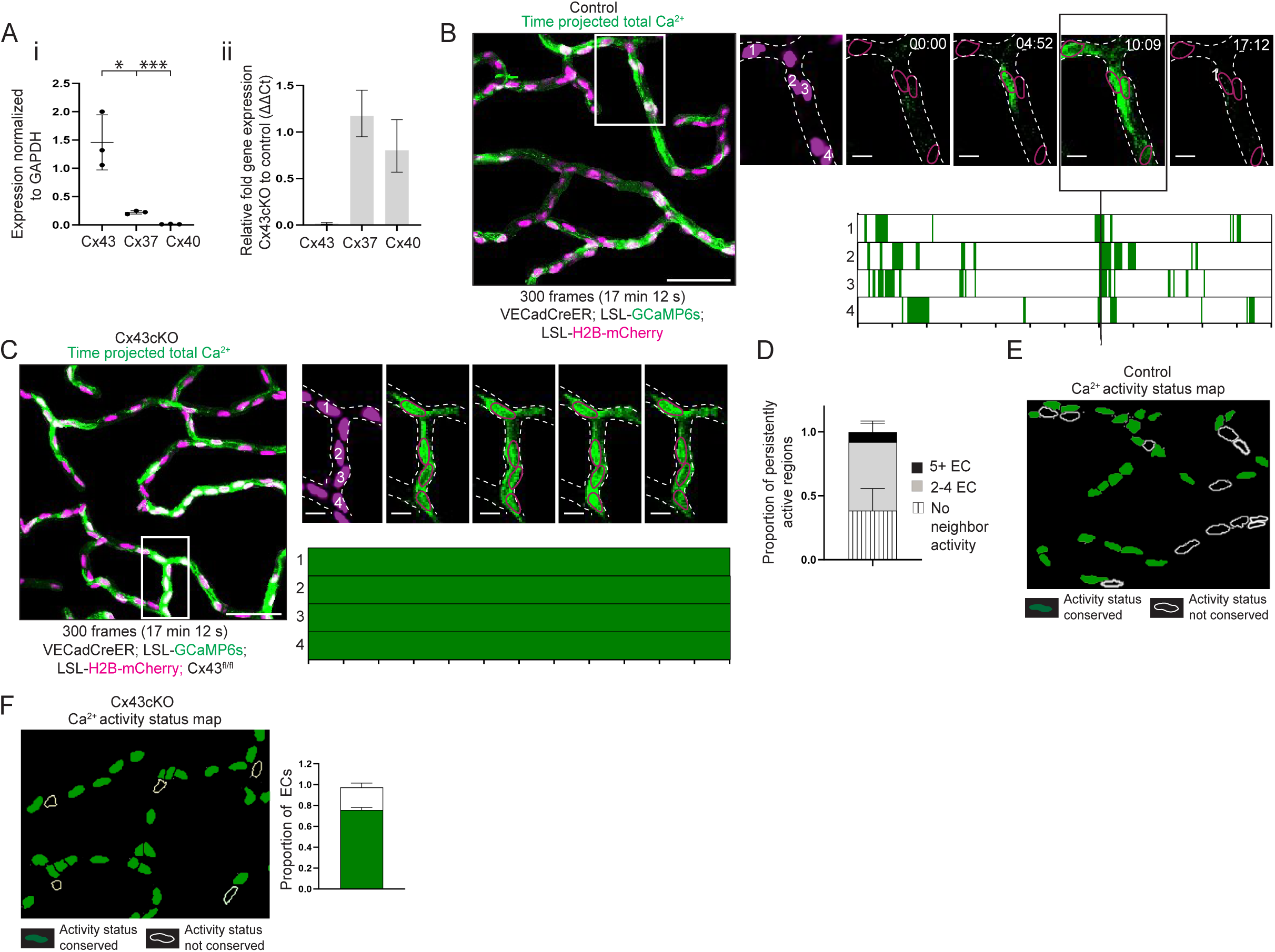
Spatial and molecular analyses of Ca^2+^ dynamics across control and Cx43cKO mice (A) **(i)** Expression of vascular connexins (Cx43, Cx37, and Cx40) in sorted skin ECs, normalized to GAPDH expression. *n=* 3 mice. **(ii)** Relative expression (ΔΔC_t_ method) of vascular connexins after Cx43cKO, compared to control mice. Connexin expression in Cx43cKO mice is first normalized to GAPDH, and then to expression in control mice. *n=* 3 Cx43cKO 3 control mice. (**B)** *Left*: Max intensity projection of GCaMP6s signal (green) with H2B-mCherry signal (magenta) from 300 frame (17 minutes 12 seconds) recording of skin capillary ECs in control mice (scale bar: 50 µm). *Right*: Inset of a region with Ca^2+^ activity occurring simultaneously across 4 ECs (numbered and drawn in magenta outline matching H2B-mCherry) (scale bar: 10 µm). *Bottom*: Ca^2+^ events and their durations over recording time for each EC. Black line across plots indicates the timepoint when all 4 ECs simultaneously display Ca^2+^ activity. **(C)** *Left*: GCaMP6s and H2B-mCherry signal from skin capillary ECs in Cx43cKO mice (scale bar: 50 µm). *Right*: Inset of a region with Ca^2+^ activity occurring simultaneously across 4 persistently active ECs (scale bar: 10 µm). *Bottom*: Ca^2+^ events for each persistently active EC. **(D)** Proportion of persistently active regions involving different EC cluster sizes. *n* = 10 regions from 3 mice. **(E)** Ca^2+^ activity status map for control ECs revisited after 14 days. Non-conserved activity status (white outline) and conserved activity status (green). *n=* 4 regions from 3 mice. **(F)** Ca^2+^ activity status map for Cx43cKO ECs revisited after 14 days. Proportion of ECs by their conservation of activity status. Chi-square analysis P<0.0001. *n=* 7 regions from 4 mice.

**Figure S4.**
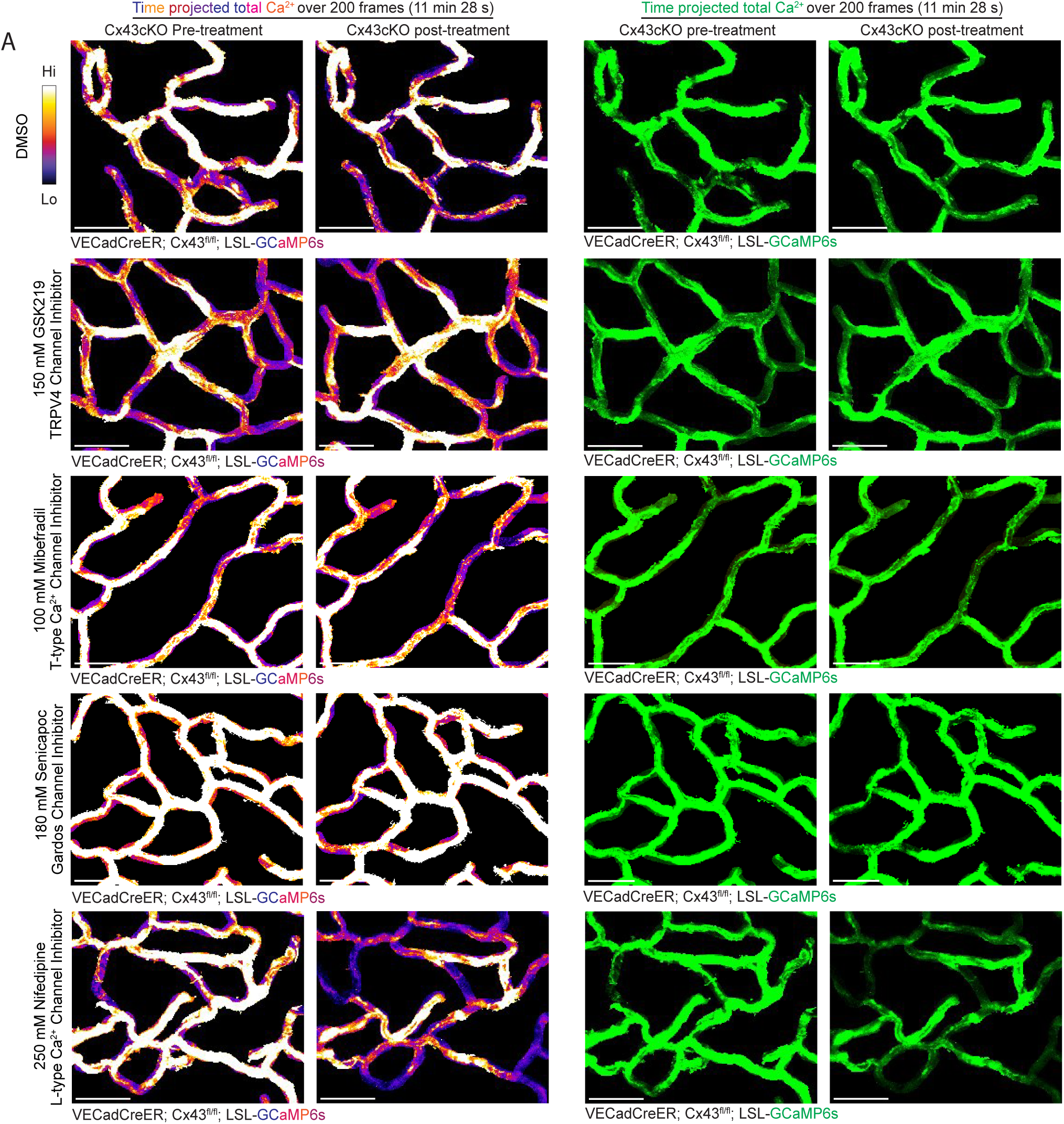
L-type VGCC inhibition, relative to inhibiting T-type VGCCs, KCa3.1 channels, and TRPV4, decreases Ca^2+^ activity after loss of Cx43. **(A)** Max intensity projection of GCaMP6s signal in green and fire lookup table from 200 frame (11 minutes 28 seconds) recording of skin capillary ECs in Cx43cKO mice before and after treatment with DMSO, GSK219, Mibefradil, Senicapoc, and Nifedipine. Fire lookup table allows for easier visualization of changes in Ca^2+^ signaling intensity. *n=* 3 mice for each condition (scale bar: 50 µm).

**Figure S5.**
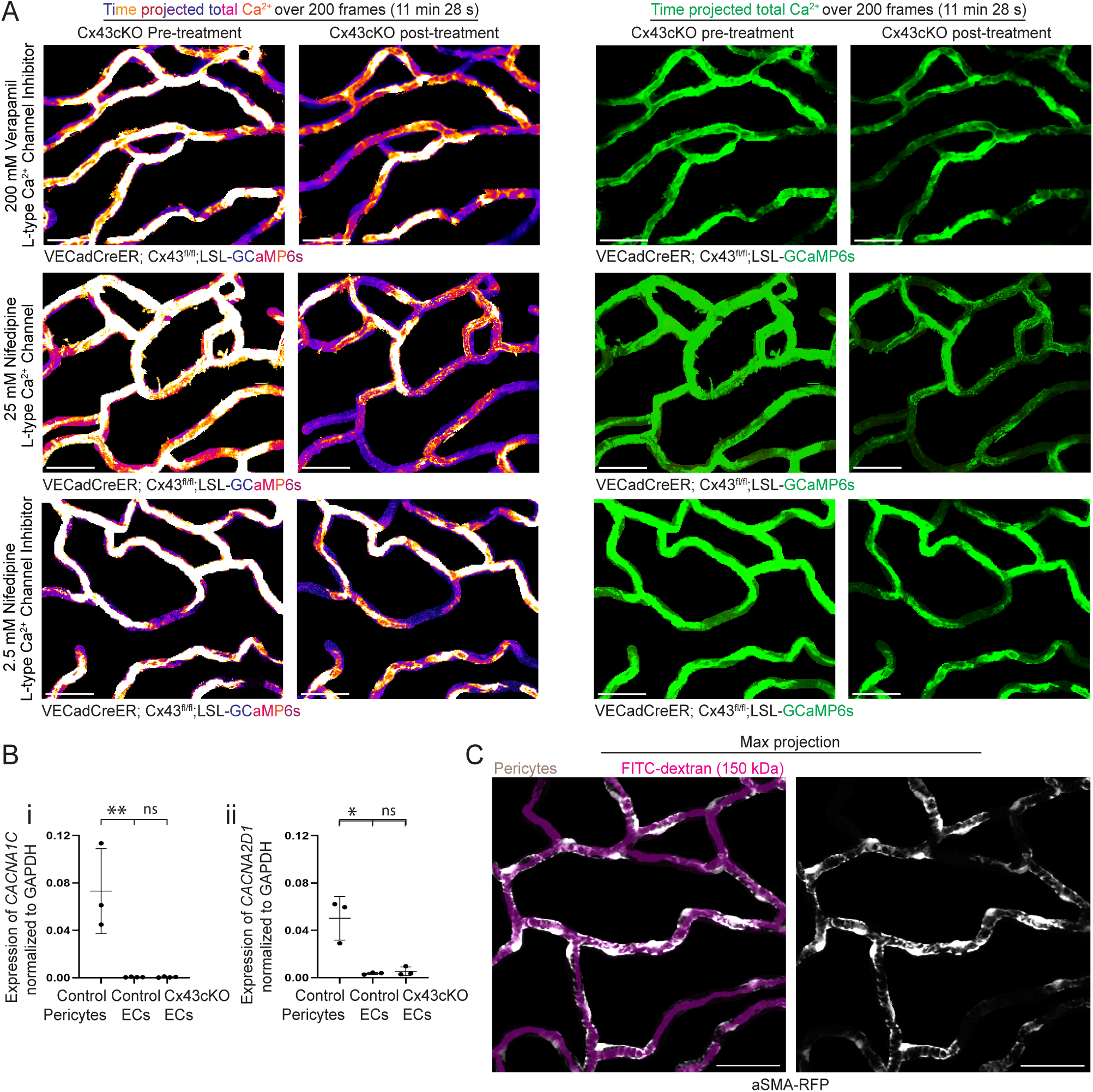
Inhibition of L-type VGCCs specifically decreases Ca^2+^ activity after loss of Cx43 through non-cell autonomous regulation. **(A)** Max intensity projection of recording before and after treatment with verapamil, and nifedipine (10 times and 100 times lower dose). Fire lookup table allows for easier visualization of changes in Ca^2+^ signaling intensity. *n=* 3 mice for each condition (scale bar: 50 µm). **(B) (i)** Expression of *CACNA1C* in sorted skin ECs and smooth muscle actin (SMA+) pericytes during homeostasis, and skin ECs after loss of Cx43, normalized to GAPDH expression. P=0.009 and ns: P>0.05 unpaired t-tests respectively; *n=* 3 aSMA-RFP mice to isolate SMA+ pericytes, n=4 control and Cx43cKO mice. **(ii)** Expression of *CACNA2D1* normalized to GAPDH expression. P=0.0479 and ns: P>0.05 unpaired t-tests respectively; *n=* 3 aSMA-RFP mice, n=4 control and Cx43cKO mice. **(C)** Representative images of capillary region (grey) in aSMA-RFP mice with 150 kDa FITC dextran (magenta) (scale bar: 50 µm).

**Figure S6.**
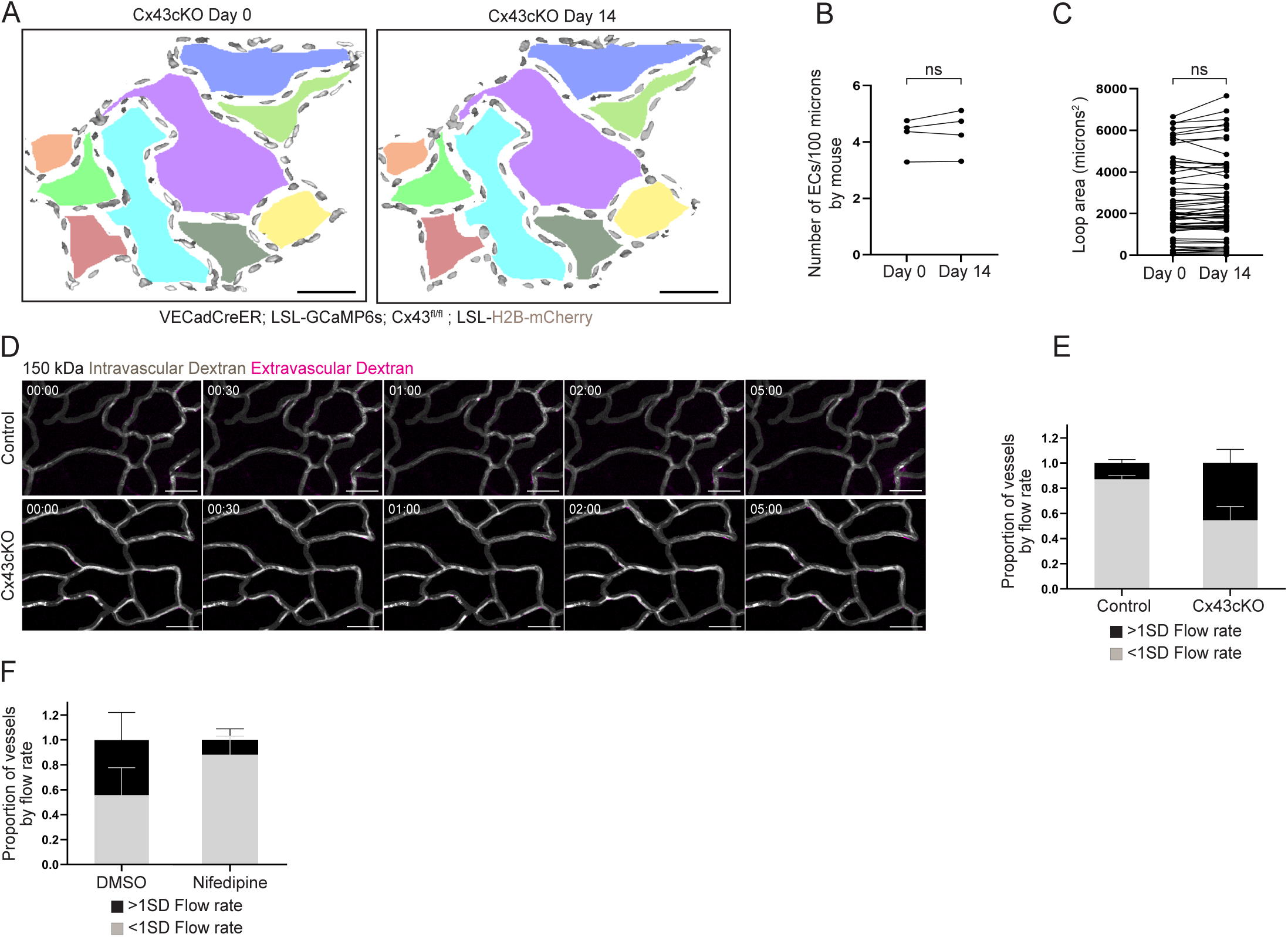
Architectural, barrier, and flow functional analyses downstream of Ca^2+^ elevation after loss of Cx43. **(A)** *Left:* Max intensity projection of capillary ECs in a Cx43cKO mouse with H2B-mCherry signal (gray) and color-coded architectural loops (scale bar: 50 µm). *Right:* Max intensity projection of the same cells revisited after 14 days, with the same colors corresponding to the same vessel loops (scale bar: 50 µm). **(B)** Number of ECs per 100 µm of the same regions revisited on Day 0 and Day 14 in Cx43cKO mice; ns: P>0.05, paired t-test, *n=*6 regions total from 4 mice. **(C)** Area of the same architectural loops revisited on Day 0 and Day 14 in Cx43cKO mice; ns: P>0.05, paired t-test, *n=*59 total vessel loops from 4 mice. **(D)** Representative single time point images of 150 kDa intravascular dextran (grey) with no extravascular dextran in magenta (scale bar: 50 µm). *n=* 3 mice each for control and Cx43cKO. **(E)** Proportion of vessels by their flow rate in control and Cx43ckO mice, separated by >1SD fast flow vessels (black) and <1SD flow vessels (grey). *n=*45 vessels from 3 mice for each group. **(F)** Proportion of vessels by their flow rate in Cx43cKO mice treated with DMSO or nifedipine. *n=*45 vessels from 3 mice for each group.

**Figure S7.**
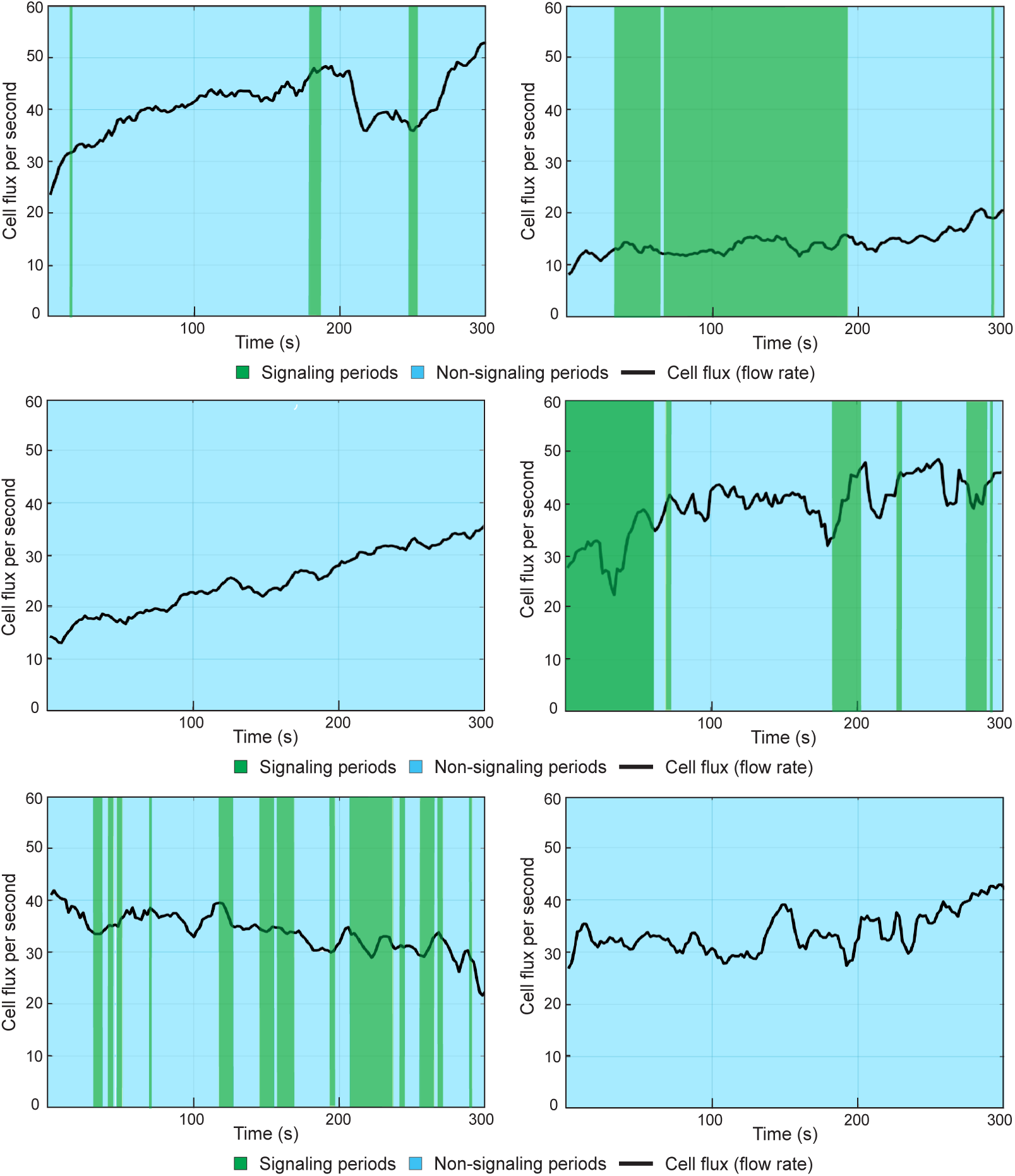
Vessel flow and Ca^2+^ signaling during homeostasis. **(A)** Example graphs across multiple vessels representing cell flux per second or flow rate during line scanning over a 5 min period. Flow rate (black line) is represented over signaling (green) and non-signaling (light blue) time periods.

**Movie 1 EC Ca^2+^ activity in skin capillaries is widespread and heterogenous during homeostasis** Timelapse of GCaMP6s signal (green) and H2B-mCherry (magenta) from 300 frame (17 minutes 12 seconds) recording of skin capillary ECs, imaged at a frame rate of 3.44 s/frame (scale bar: 20 µm). Timelapse is followed by a max intensity projection of GCaMP6s signal represented with fire lookup table. Color scale indicates GCaMP6s signal over recording time.

**Movie 2 EC Ca^2+^ activity status is conserved on a single-cell level after 24 hours** *Left*: Timelapse of GCaMP6s signal (green) and H2B-mCherry (magenta) from 300 frame (17 minutes 12 seconds) recording of skin capillary ECs on baseline Day 0, imaged at a frame rate of 3.44 s/frame (scale bar: 20 µm). *Right:* Timelapse of region from Day 0 revisited after 24 hours. Timelapse is followed by a max intensity projection of GCaMP6s signal for Day 0 and Day 1 represented with fire lookup table. Color scale indicates GCaMP6s signal over recording time.

**Movie 3 EC Ca^2+^ activity status is conserved on a single-cell level after 14 days** *Left*: Timelapse of GCaMP6s signal (green) and H2B-mCherry (magenta) from 300 frame (17 minutes 12 seconds) recording of skin capillary ECs on baseline Day 0, imaged at a frame rate of 3.44 s/frame (scale bar: 20 µm). *Right:* Timelapse of region from Day 0 revisited after 2 weeks. Timelapse is followed by a max intensity projection of GCaMP6s signal for Day 0 and Day 14 represented with fire lookup table. Color scale indicates GCaMP6s signal over recording time.

**Movie 4 Cx43cKO leads to sustained EC Ca^2+^ activity** *Left*: Timelapse of GCaMP6s signal (green) and H2B-mCherry (magenta) from 300 frame (17 minutes 12 seconds) recording of skin capillary ECs from control mice, imaged at a frame rate of 3.44 s/frame (scale bar: 20 µm). *Right:* Timelapse of capillary ECs from Cx43cKO mice. Timelapse is followed by a max intensity projection of GCaMP6s signal for control and Cx43cKO mice, represented with fire lookup table. Color scale indicates GCaMP6s signal over recording time.

**Movie 5 Cx43cKO leads to increase in persistently active ECs after 2 weeks** *Left*: Timelapse of GCaMP6s signal (green) and H2B-mCherry (magenta) from 300 frame (17 minutes 12 seconds) recording of skin capillary ECs from Cx43cKO mice on baseline Day 0, imaged at a frame rate of 3.44 s/frame (scale bar: 20 µm). *Right:* Timelapse of region from Day 0 revisited after 2 weeks, on Day 14. Timelapse is followed by an average intensity projection of GCaMP6s signal for Cx43cKO mice on Day 0 and Day 14, represented with fire lookup table. Average intensity projections allow better visualization of persistently active regions in white. Color scale indicates GCaMP6s signal over recording time.

**Movie 6 L-type VGCC inhibition does not affect EC Ca^2+^ activity in control mice** *Top Left*: Timelapse of GCaMP6s signal (green) and H2B-mCherry (magenta) from 200 frame (11 minutes 28 seconds) recording of skin capillary ECs from control mice prior to DMSO treatment, imaged at a frame rate of 3.44 s/frame (scale bar: 20 µm). *Top Right*: Timelapse of revisited region after DMSO treatment. *Bottom Left*: Timelapse of ECs prior to nifedipine treatment. *Bottom Right*: Timelapse of revisited region after nifedipine treatment. All timelapses are followed by max intensity projection of GCaMP6s signal represented with fire lookup table. Color scale indicates GCaMP6s signal over recording time.

**Movie 7 L-type VGCC inhibition decreases EC Ca^2+^ activity after Cx43cKO** *Top Left*: Timelapse of GCaMP6s signal (green) and H2B-mCherry (magenta) from 200 frame (11 minutes 28 seconds) recording of skin capillary ECs from Cx43cKO mice prior to DMSO treatment, imaged at a frame rate of 3.44 s/frame (scale bar: 20 µm). *Top Right*: Timelapse of revisited region after DMSO treatment. *Bottom Left*: Timelapse of ECs prior to nifedipine treatment. *Bottom Right*: Timelapse of revisited region after nifedipine treatment. All timelapses are followed by max intensity projection of GCaMP6s signal represented with fire lookup table. Color scale indicates GCaMP6s signal over recording time.

